# Stochastic survival of the densest and mitochondrial DNA clonal expansion in ageing

**DOI:** 10.1101/2020.09.01.277137

**Authors:** Ferdinando Insalata, Hanne Hoitzing, Juvid Aryaman, Nick S. Jones

## Abstract

The expansion of deleted mitochondrial DNA molecules has been associated with ageing ^1,2^, particularly in skeletal muscle fibres ^3–5^; its mechanism has remained unclear for three decades. Previous accounts have assigned a replicative advantage to the deletions ^6–8^, but there is evidence that cells can, instead, selectively remove defective mitochondrial DNA ^9^. Here we present a spatial model that, without a replicative advantage, but instead through a combination of enhanced density for mutants and noise, produces a wave of expanding mutations with speeds consistent with experimental data ^10^. A standard model based on replicative advantage yields waves that are too fast. We provide a formula that predicts that wave-speed drops with copy number, consonant with experimental data. Crucially, our model yields travelling waves of mutants even if mutants are preferentially eliminated. Additionally, we predict that experimentally observed mutant loads can be produced by *de novo* mutation rates that are drastically lower than previously thought for neutral models ^11^. Given this exemplar of how noise, density and spatial structure affect muscle ageing, we introduce the mechanism of stochastic survival of the densest, an alternative to replicative advantage, that may underpin other evolutionary phenomena.

## Introduction

The accumulation of mitochondrial DNA (mtDNA) mutations to high levels has been repeatedly linked to ageing ^1,2^, especially in postmitotic tissues such as neurons or muscles ^12^. In these tissues, a bioenergetic defect can be triggered when the proportion of mutant mtDNA in a region exceeds a threshold value ^13,14^. Sarcopenia, the loss of skeletal muscle mass and strength with age, is widely associated with high levels of mtDNA deletions (Fig. 1A)^3–5^. A defining feature of the expansion of mtDNA deletions in muscle fibres is clonality: damaged regions of the muscles are taken over by a single deletion. The mechanism behind this phenomenon has remained unclear despite numerous authors using mathematical modelling to probe it^6–8,11,15–18^. Some models reproduce the clonal expansion by assigning a replicative advantage to mtDNA deletions ^6–8,17,18^. However, there is no definitive biological mechanism to justify the supposed replicative advantage of deletions; by contrast, there is evidence that deleted mtDNA is preferentially eliminated^9,19^. Neutral stochastic models have also been proposed ^11,15,16^, describing the clonal expansion in terms of neutral stochastic drift, but they require excessively high *de novo* mutation rates to reproduce observed mutant loads in short-lived animals ^17,20^. Moreover, most existing models neglect the spatial structure of muscle fibres. Another widely reported feature of the clonal expansion is the higher density of deletions: regions of the muscle fibre taken over by the mutations present an approximate fivefold increase^3,4,21–23^ in absolute mtDNA copy number. Molecular mechanisms to account for increased mitochondrial density have been previously proposed, including the “maintenance of wildtype” hypothesis ^15,24^, and homeostasis on ATP production ^25,26^, proteome status ^16,27,28^ or mtDNA copy number^28^.

**Figure 1:**
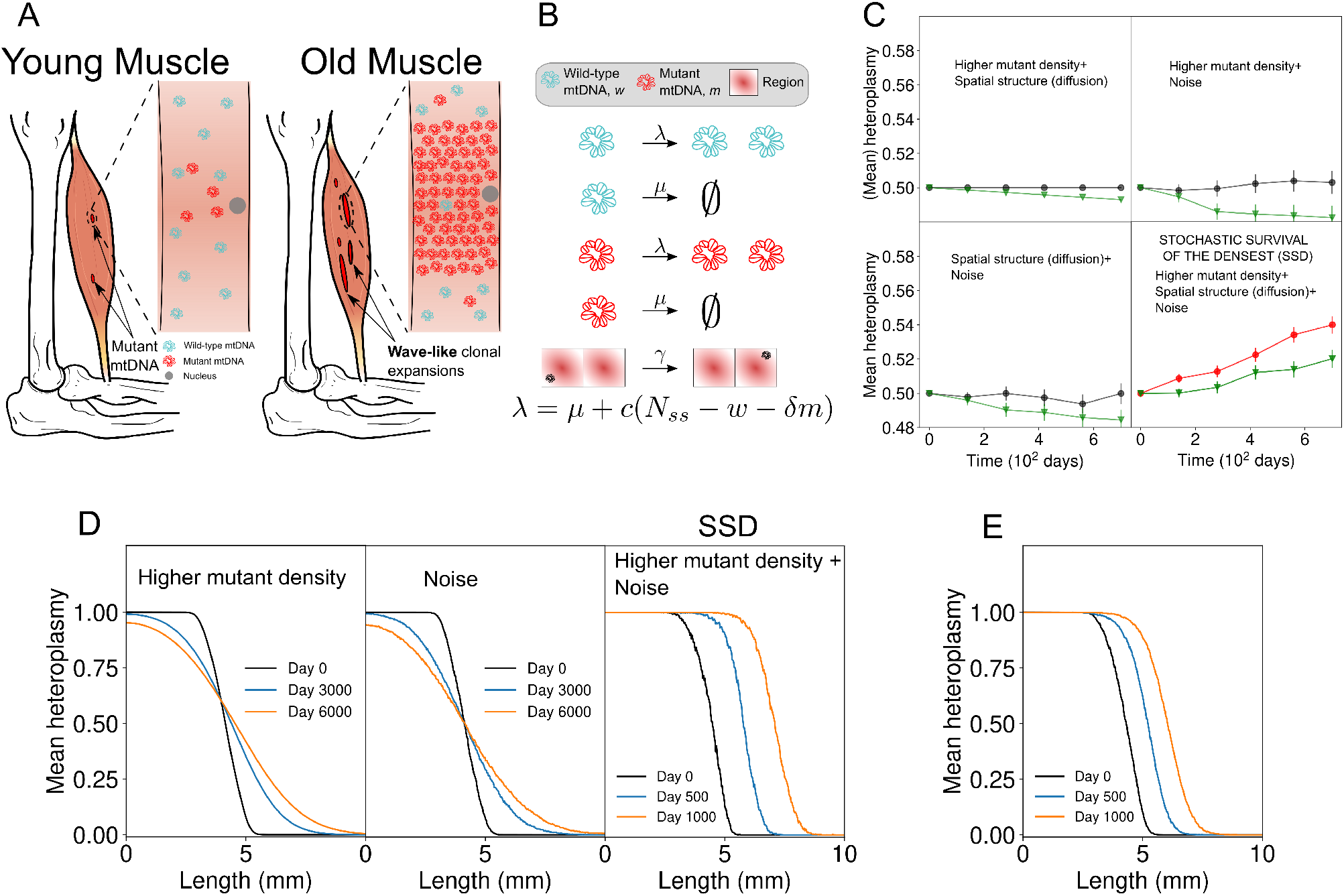
Stochastic survival of the densest (SSD) can produce increases in the proportion of mutants even if they are subject to higher degradation rates than wildtypes. (A) Dysfunctional mtDNA mutants expand in muscle fibres with age in a wave-like manner, leading to defects in OXPHOS. (B) In a spatially extended system, the possible events are birth and death of a wildtype (first two), birth and death of a mutant (third and fourth), a mutant or wildtype hops to a neighboring unit (fifth). Births happen at rate *λ*(*w, m*) (defined at the bottom), deaths at constant rate *μ*, hopping at constant rate *γ*. (C) SSD (bottom right subpanel) is observed in the presence of noise, spatial structure (with diffusion) and higher density of mutants, which lead to increase in mean heteroplasmy in a neutral model (red line) and even with higher degradation of mutants (green line). If any of the factors is missing, as in the other subpanels, heteroplasmy stays constant in a neutral model (black) or decreases with preferential elimination of mutants (green). Error bars are standard error of the mean. (D) In a spatially structured model, SSD drives a travelling wave of mutants only in the presence of noise and higher mutant density (rightmost subpanel), while the high-heteroplasmy front diffuses away if either noise (left) or higher mutant density (middle) are missing. (E) For the same model as in D, if mutants are preferentially degraded an invasive wave can still occur, because of SSD. If any of the three factors of SSD is missing, mutants subject to a higher degradation rate go extinct instead (e.g. Fig. S4B).

Here we introduce a stochastic model of the evolution of mtDNA in skeletal muscle fibres that, consonant with data, predicts the clonal expansion of deletions without assuming a replicative advantage, even allowing mutants to be subject to preferential elimination, and drastically lowers the *de novo* mutation rate required to account for a given mutant load. The model’s parameters have all a clear biological interpretation and are estimated from published experimental data. The wave of mutants we observe requires the model’s stochasticity and is an effect which promises wider evolutionary application.

## Clonal waves of mutants

The building block of our approach is a simple stochastic model that describes a population of mtDNA, wildtype *w* and mutants *m*, evolving under central regulation by the nucleus. The main quantity of interest is heteroplasmy *h*: the proportion of the population that is mutant. Importantly, we principally focus on the mechanism by which a pre-existing mutation reaches high heteroplasmy through clonal expansion; we will later consider *de novo* mutations occurring continuously through time. The model is neutral: the two species have an identical constant degradation rate *μ*, and an identical replication rate *λ* given by

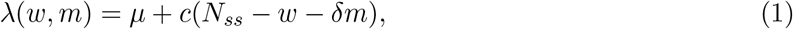

with parameters *c, N_ss_, δ.* The replication rate is scaled by the distance of a count, *w* + *δm*, of the current population size from a target population *N_ss_*. Crucially, mutants contribute less to the current population than wildtypes, the relative size of this contribution being the parameter 0 ≤ *δ* < 1. The parameter *c* quantifies the strength of control exerted by the nucleus (SI.1.1). In SI.1 we show that 0 ≤ *δ* < 1 means mutants are the denser species. The focus of this work is providing a mechanistic model of the clonal expansion of the mtDNA deletions, and we can be agnostic to the detailed molecular mechanism behind their observed higher density^3,4,21–23^.

The single-unit model is summarised by the first four reactions in Fig. 1B. In this setting, for 0 ≤ *δ* < 1, mean heteroplasmy stays constant for both the deterministic (Eq. (S4)) and stochastic (Eq. (S16)) versions of this system. Although mean heteroplasmy is constant, mean mutant copy number can increase through stochastic mechanisms (Eq. S14)^15,29,30^. Coupling two of these units together, allowing exchange of mtDNA molecules between them at a constant per-capita rate *γ* (last reaction Fig. 1B), makes no difference under deterministic dynamics (Fig. 1C, top left, black), makes no difference under stochastic dynamics if *δ* = 1 (Fig. 1C, bottom left, black) but, we observe an increase in mean heteroplasmy for stochastic dynamics with diffusion and 0 ≤ *δ* < 1 (Fig. 1C, bottom right, red). Fig. 1C illustrates that this effect requires 1) spatial structure with diffusion of molecules between units, 2) stochasticity, 3) higher density of mutants (0 < *δ* < 1). For this reason, we term this novel mechanism stochastic survival of the densest (SSD). In this paper we link SSD to the clonal expansion of deletions in skeletal muscle fibres. We model a fibre as a chain of units, each containing an mtDNA population evolving under nuclear control, and exchanging mtDNA molecules between neighbours.

In our spatial model of skeletal muscle fibres – a chain of units with diffusion of mtDNA molecules – the high-heteroplasmy front diffuses away without noise (Fig. 1D, left) or without higher mutant density (Fig. 1D, middle) and only advances in the presence of noise and higher density mutants (Fig. 1D, right). The wave of advance of mutants requires spatial structure, noise and higher mutant density. When these three elements are present, SSD predicts the wave-like expansion of mutants even if they are preferentially degraded (Fig. 1E), whereas in a deterministic model a higher degradation rate for mutants leads to their extinction (Fig. S4B).

## SSD matches clonal expansion in muscles

Experimental data (Fig. 2A, ^21^), shows the characteristic spatial profile of heteroplasmy data, with high-heteroplasmy (OXPHOS-defective) regions flanked by transition regions to low or zero heteroplasmy. This heteroplasmy profile is well described by a sigmoid (SI.16), the shape expected for a travelling wave (SI.14). This suggests that the expansion is a wave-like phenomenon. We have estimated the speed of this wave by analysing data from Ref. [10] on the length of abnormal regions in rhesus monkeys and age of the subject. Regressing the lengths against age (Fig. 2B), we observe a relationship (*p* = 5 · 10^−4^) which is approximately linear (*R*^2^ = 0.76) and corresponds to an average wave-speed of (0.131 ± 0.025)μm/day. Fig. 2C, relative to the same fibre as in 2A^21^, highlights the key fact that the absolute copy number in high-heteroplasmy regions is larger than in normal regions: mutants are the denser species. This is an established phenomenon known for over 20 years ^3,4,15,22,23^. We find that a standard model which assigns a replicative advantage to mutants, and accounts for the wavefront’s elevation in copy number (SI.9), predicts a wave-like expansion of ≈ 40 μm/day (Fig. 2D), 300 times faster than the observed speed. In contrast, SSD predicts a wave-like expansion of deletions with a speed of ≈ 0.2 μm/day (Fig. 2 E). We have estimated the parameter values for both models from data in the literature (see SI.9).

**Figure 2:**
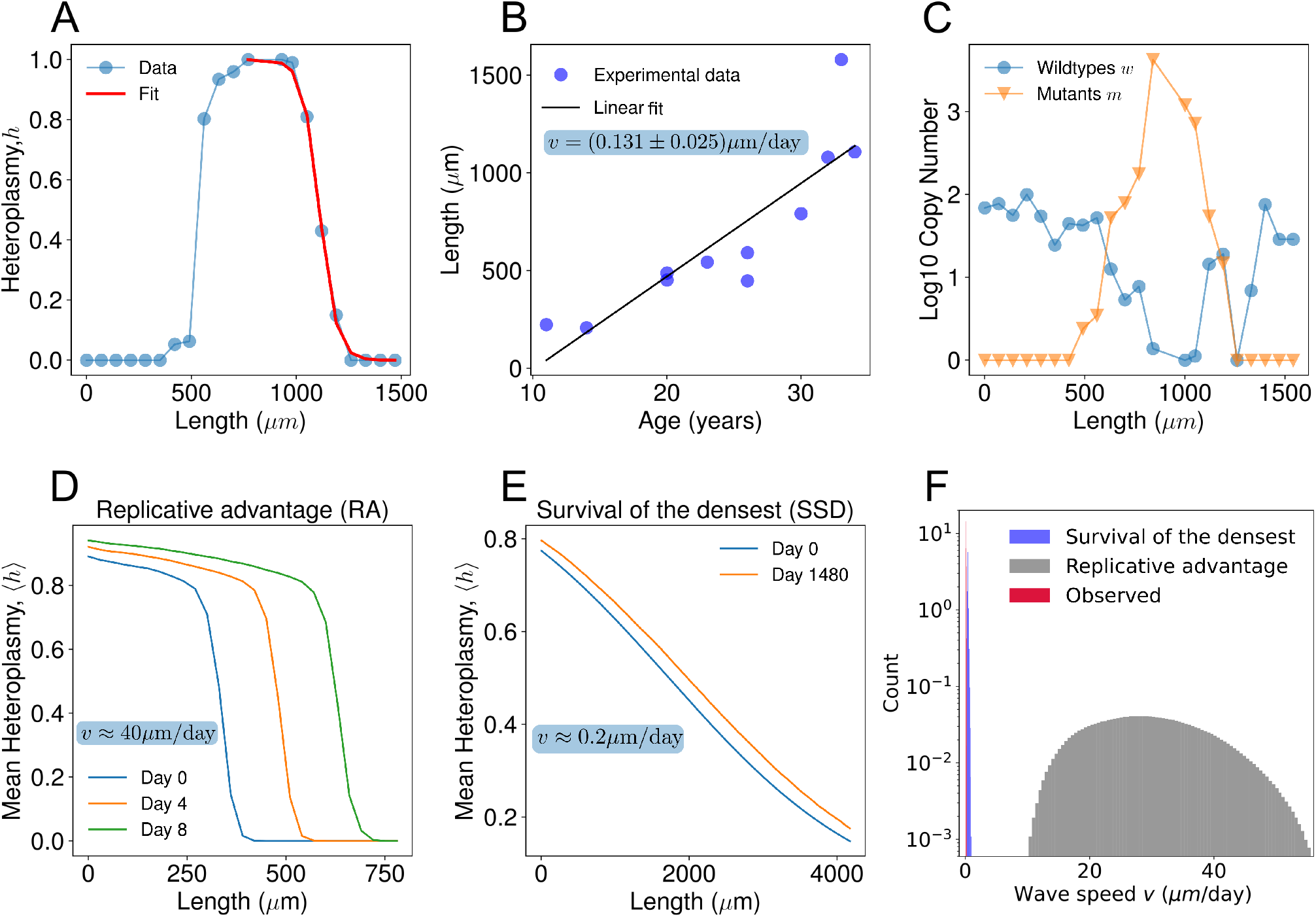
Stochastic survival of the densest predicts a wave-like expansion of mtDNA mutants at a speed in agreement with experimental observations, while a standard replicative advantage model predicts a speed a factor of ≈ 300 too large. A) Spatial profile of mutant fraction (heteroplasmy) along a human skeletal muscle fibre ^21^. The heteroplasmy profile follows a sigmoid (SI.16), the shape expected for a travelling wave. (B) Experimental data on the length of abnormal regions of muscle fibres against age in rhesus monkeys, from Ref. [10]. An approximate linear relationship is found (*R*^2^ = 0.76, *p* = 5 · 10^−4^), compatible with a wave-like expansion with speed (0.131 ± 0.025)μm/day (linear fit). (C) Spatial structure of copy number for wildtype (blue) and mutants (orange) for the same muscle fibre as in panel A. The heteroplasmic regions present a higher absolute copy number, i.e. mutants are present at a higher density. (D) Stochastic simulations of a spatially extended model with a replicative advantage for mutants, with our best estimate of the model parameters (see SI.9) for muscle fibres of rhesus monkeys predicts a wave-like expansion with a speed of ~ 40μm/day, 300 times faster than the observed speed. (E) Simulations of survival of the densest, with same death and replication rate for wildtypes and mutants, yields a mutant wave speed of ~ 0.2μm/day for the fibres or rhesus monkeys, which is comparable with experimental observations (see (C)). (F) Inserting probabilistic estimates of the model parameters (see SI.9) into Eq. (2), we find that survival of the densest predicts a distribution for the wave speed (red histogram) compatible with observations (blue), whereas a wave driven by replicative advantage with the same model parameters is two orders of magnitude faster (grey).

By measuring the simulated wave-speed for 110 combinations of parameters, we found that it is well described (*R*^2^ = 0.99) by the phenomenological formula:

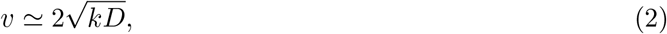

where *D* is the diffusion coefficient of mtDNA along the fibres and with 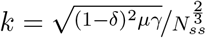

Eq. (2) is analogous to the wave-speed formula for the wave of advance of advantageous mutants introduced by Fisher and Kolmogorov^31,32^. In our case *k* can be seen as an effective selective advantage for mutants induced by SSD, in contrast with the replicative advantage (RA) driving the original Fisher-Kolmogorov waves. In light of this analogy, our model can be seen as a reaction-diffusion system where the reaction component emerges from the combined effect of noise and higher mutant density.

We obtained probability distributions for the wave-speeds predicted by SSD and a RA model, by inserting draws from the distributions of parameter values (given in SI.9) into Eq. (2), with the appropriate interpretation of *k* for the two models. The predicted distributions are plotted in Fig. 2F, together with the distribution of the experimentally observed wave-speed obtained via linear fit (Fig. 2B). After accommodating this parametric uncertainty, SSD remains much superior to RA at reproducing the observed speed.

## Linking ageing, wave-speed and copy number

As Eq. (2) states, SSD predicts a wave-speed that decreases when copy number (per nucleus) increases. Indeed, the expansion of mutants is driven by stochastic fluctuations, whose effect generally becomes smaller for larger population size (see also Eq. (S14)). In contrast, a standard model based on an RA predicts a wave-speed that increases with population size ^31,33^. This allows us to test SSD and RA models against other experimental observations. It has been found that copy number depletion caused by antiretroviral therapy^34^ or AKT2 deficiency ^35^ is associated with enhanced sarcopenia. Likewise, statins are well known for increasing the risk of sarcopenia ^36–38^ and have consistently been associated with reduction in mitochondrial copy number ^39–41^. Conversely, increase in mtDNA content through exercise ^42,43^ or overexpression of TFAM ^44^ and parkin ^45^ have been found to protect against sarcopenia and muscle atrophy.

Skeletal muscle fibres can be broadly classified into Type 1 (oxidative) and Type 2 (glycolytic) fibres ^46^. The former rely on OXPHOS to function and typically have twice as many mitochondria as the latter ^10,47,48^, that depend on glycolysis. It is known that Type 2 fibres are more affected by sarcopenia with ageing ^3,10,49–52^. Small, short-lived animals like rodents show sarcopenia on a time scale of years (from ≈ 2 years) and have a larger proportion of Type 2 fibres compared to longlived animals such as rhesus monkeys and humans ^34^, that exhibit sarcopenia on a longer time scale (decades). While there are other physiological differences between the fibre types, this is a further link between smaller mtDNA copy numbers and faster mutant expansion. The leading edge of a faster wave is flatter than that of a slower wave (e.g. Ref. [53], summarised in SI.14). By exploiting this property, it is possible to compare the speeds of two waves by examining their shapes. Data on muscle fibres for humans ^3^ (Fig. 3A, B) and rats ^4^ (Fig. 3C, D) show that flatter – hence faster – waves of mutants propagate along fibres with smaller copy numbers. All these observations support our model’s prediction of an inverse relationship between copy number and wave-speed, opposite to that predicted by a standard RA model ^33^.

**Figure 3:**
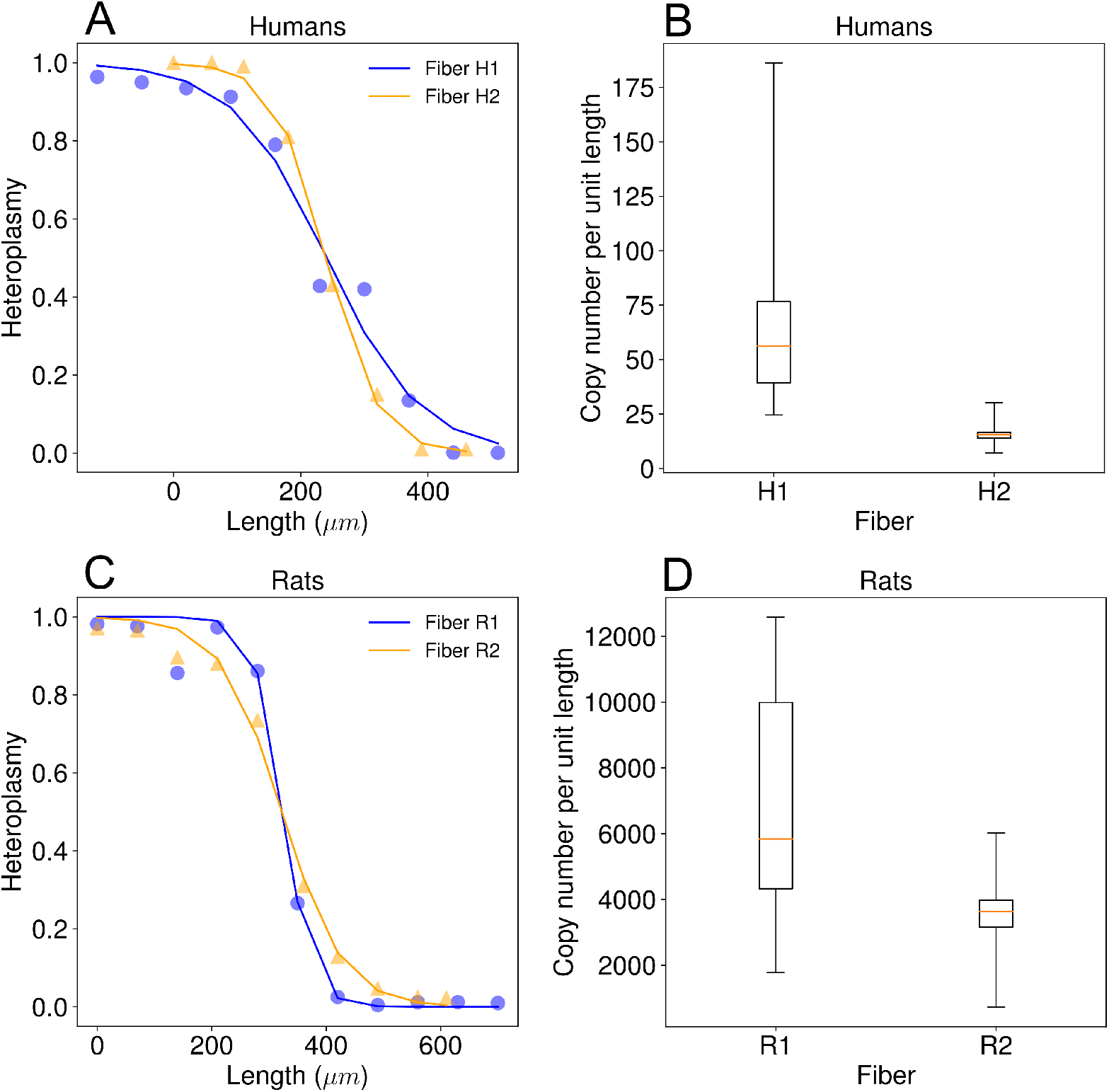
A steeper wave of mutants propagates more slowly in fibres which have a higher copy number per unit length, in agreement with the predictions of stochastic survival of the densest. (A) Significant difference in the steepness of wavefronts of two human muscle fibres H1 and H2: *τ*_*H1*_ = (2.4 ± 0.1) · 10^−2^*/*μm for H1 and *τ*_*H2*_ = (1.4 ± 0.2) · 10^−2^/μm for H2 (MLE). Data from Ref. [3]. According to the mathematics of travelling waves (SI.14), the steeper wave is slower. (B) Comparison between the corresponding *N_ss_* of the two fibres H1 and H2. We have found evidence (*p* = 10^−4^, *d* = 1.48, one-sided Welch’s *t*-test) that the average copy number per unit length (a slice of length 20 μm here) in normal regions of the two fibres H1 and H2, *N_ss_* in our model, is higher for the steeper, and hence slower, wave. This is in qualitative agreement with the predictions of our stochastic survival of the densest model that wave speed decreases with copy number. In contrast, a model based on replicative advantage predicts that speed increases with copy number. (C) Two muscle fibres in rats, R1 and R2, present waves of mutants with significantly different steepness of the waveform: *τ*_*R1*_ = (4.0 ± 0.6) · 10^−2^ / μm for R1 and *τ*_*R*2_ = (1.8 ± 0.2) · 10^−2^ /μm for R2 (MLE). (D) We have found an indication (*p* = 0.06, *d* = 1.00, one-sided Welch’s *t*-test) that the average copy number per unit length in normal regions of the two fibres R1 and R2, is higher for the steeper wave (R1). Data from Ref. [4].

Importantly, the travelling wave of mutants with inverse relationship between speed and copy number is observed not only in the case of linear feedback control (Eq. (1)), but also for other controls encoding a higher density of mutants (see SI.13), provided that stochasticity and spatial structure (with diffusion) are present.

## Low mutation rates can yield large mutant load

Previous neutral models of mtDNA dynamics in skeletal muscle fibres require high *de novo* mutation rates *R_mut_* to explain the observed mutational loads in fibres of short-lived animals ^4,17,20,54^. In turn, these high mutation rates produce an unrealistically high mutational diversity, at odds with the observed clonality of the expansion of deletions: this shortcoming has motivated theorists to develop RA models. However, previous studies modelled skeletal muscle fibres as an unstructured bulk of well-mixed mtDNAs^6,11,17^. Spatial structure is one of the three key factors in SSD (Fig. 1C, D): non-spatial models do not predict the wave-like expansion of mutations that we highlight here. Crucially, a travelling wave of mutants in a stochastic model yields a higher probability of fixation for mutants, which implies that a lower mutation rate is needed to produce a given mutant load. In Fig. 4 we plot the reciprocal of the probability that a founder mutation takes over a muscle fibre against *N_ss_* over three orders of magnitude. The inferred probability for *N_ss_* = 3500 (humans) is such that the *de novo* mutation rate required to reproduce observed mutant loads is in the range *R_mut_* = 4.1 · 10^−9^ 1.6 · 10^−7^ per replication for a typical fibre (SI.11), in line with conservative experimental estimates ^11,55^ and about three orders of magnitude smaller than estimated in the reference study ^11^. In conclusion, SSD can reproduce the observed mutant loads in skeletal muscle fibres requiring drastically lower *de novo* mutation rates than previous neutral models.

**Figure 4:**
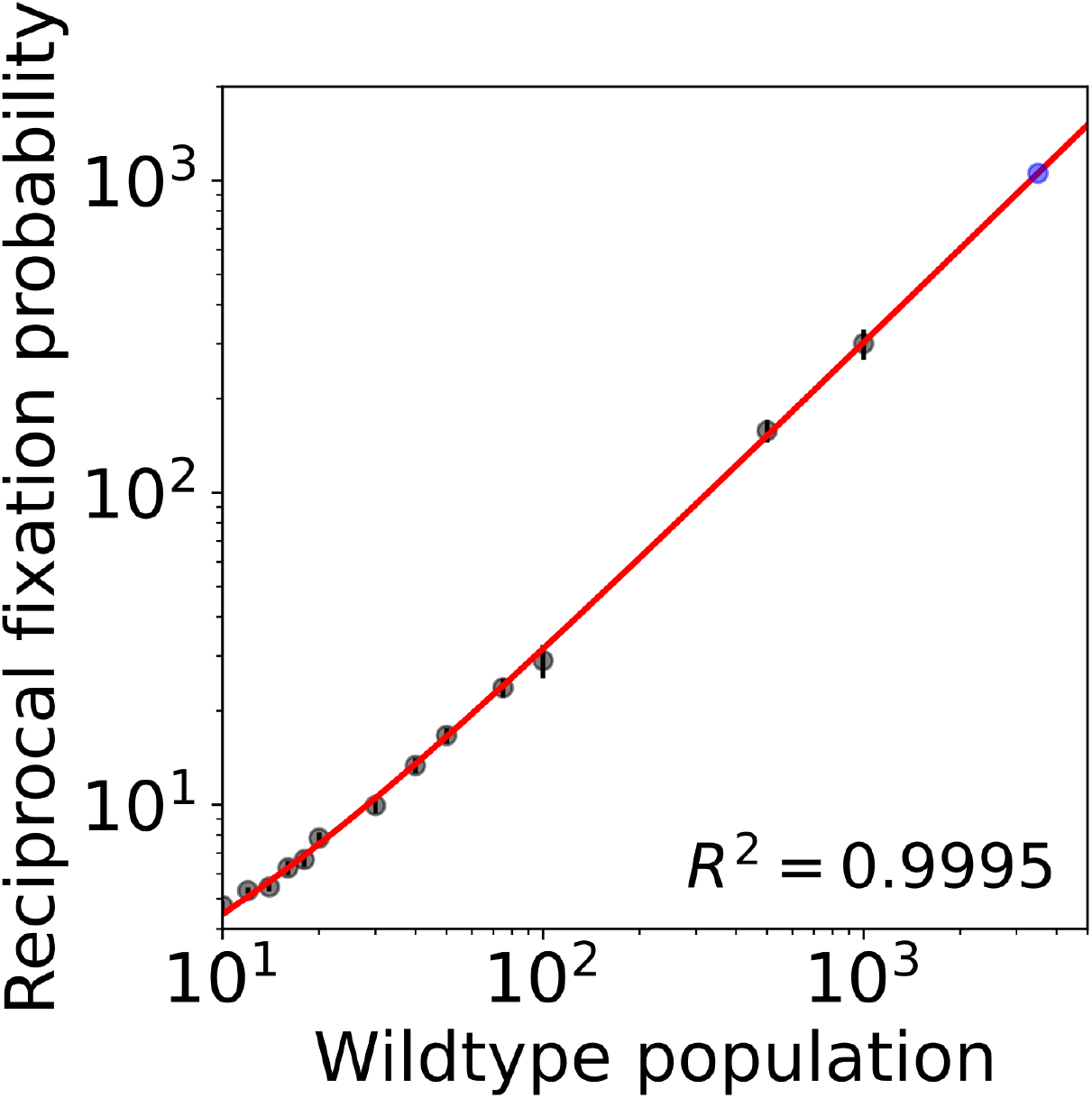
Fixation probability of a founder mutation in a muscle fibre depends on local numbers of mtDNA per nucleus (N_ss_ in our model) and is independent of the total number of mtDNA in the fibre. The reciprocal of the fixation probability *P_f_* increases linearly with *N_ss_* (see SI.11), and hence *P_f_* ≈ *α/N_ss_*, with *α* = (3.316 ± 0.002) for *N_ss_* ≫ 1. The black dots represent estimates of this probability for 10 < *N_ss_* < 1000, obtained via stochastic simulations (details in SI.12, error bars are standard deviations). For *N_ss_* = 3500, a plausible value for humans, the predicted fixation probability is *P_f_* = (9.47 ± 0.06) · 10^−4^ (blue dot).

## Discussion

We have presented a bottom-up, physically interpretable spatial stochastic model that predicts the wave-like clonal expansion of mitochondrial deletions in skeletal muscle fibres even if they are subject to preferential elimination. This counterintuitive result (stochastic survival of the densest) depends on the increased density of deletions – a widely observed fact ^3,4,21–23^ – and on the spatial and stochastic nature of the model. Previously, the clonal expansion of mitochondrial deletions has been modelled via a replicative advantage ^6–8,17,18^. We have discovered that a literature-parameterised model of this type produces a wave of deletions that is 300 times faster than observed speeds, whereas our model predicts a speed that is of the same order of magnitude as observations. We have provided a phenomenological formula for the wave-speed that has implications for therapy, since existing drugs allow us to modulate some of the parameters influencing the propagation of mutants. We have corroborated our prediction of a wave-speed decreasing with copy number, by examining how copy number changes the steepness of the wavefront and likelihood of developing sarcopenia. Finally, we have shown that the wave-like spread lowers the *de novo* mutation rate needed to reproduce the observed mutant loads by four orders of magnitude in humans. Our core claim is that the expansion of mutants is not driven by a replicative advantage, nor by neutral stochastic drift. Rather, mutants have an effective selective advantage induced by the combined effect of noise, higher density and spatial structure with diffusion (Fig. 1C, D). To our knowledge, the expansion of mtDNA deletions in skeletal muscle fibres is the first experimental candidate for what we have termed *stochastic survival of the densest*, a novel mechanism that might be the driving force behind other counterintuitive evolutionary phenomena. In the supplement, beyond giving a wider positioning of this work in the evolutionary literature (SI.6–7), we show that in our model the replication rate of all individuals increases with the proportion of mutants (SI.1.2). Mutants’ higher degradation rate might be seen as the cost of bringing this benefit (as in Refs. [30, 33, 56, 57]). An altruist can be defined as an individual that benefits others at a cost to itself ^56,58^, and there is thus a link between the mutants in our model and a specific definition of altruism. A classic setting for the wave-like spread of a trait is in the uptake of agriculture ^59^, which might not impart an explicit replicative advantage and might lead to higher death rates^60^, but nonetheless spreads, possibly due to an increase in carrying capacity of the land. We believe that this study and the simplicity of our microscopic model might pave the way for an increased recognition of this intriguing mechanism in evolutionary biology.

## Supporting information

Mathematica notebook for stochastic dimensionality reduction.

## Author contributions

## Acknowledgements

The authors acknowledge Judd Aiken and Allen Herbst for advising on skeletal muscle fibres data, Sam Greenbury and Iain Johnston for useful comments about the manuscript. NJ acknowledges the Leverhulme Trust RPG-2019-408 and EPSRC EP/N014529/1. FI is supported by Imperial College’s President’s PhD Scholarship.

---

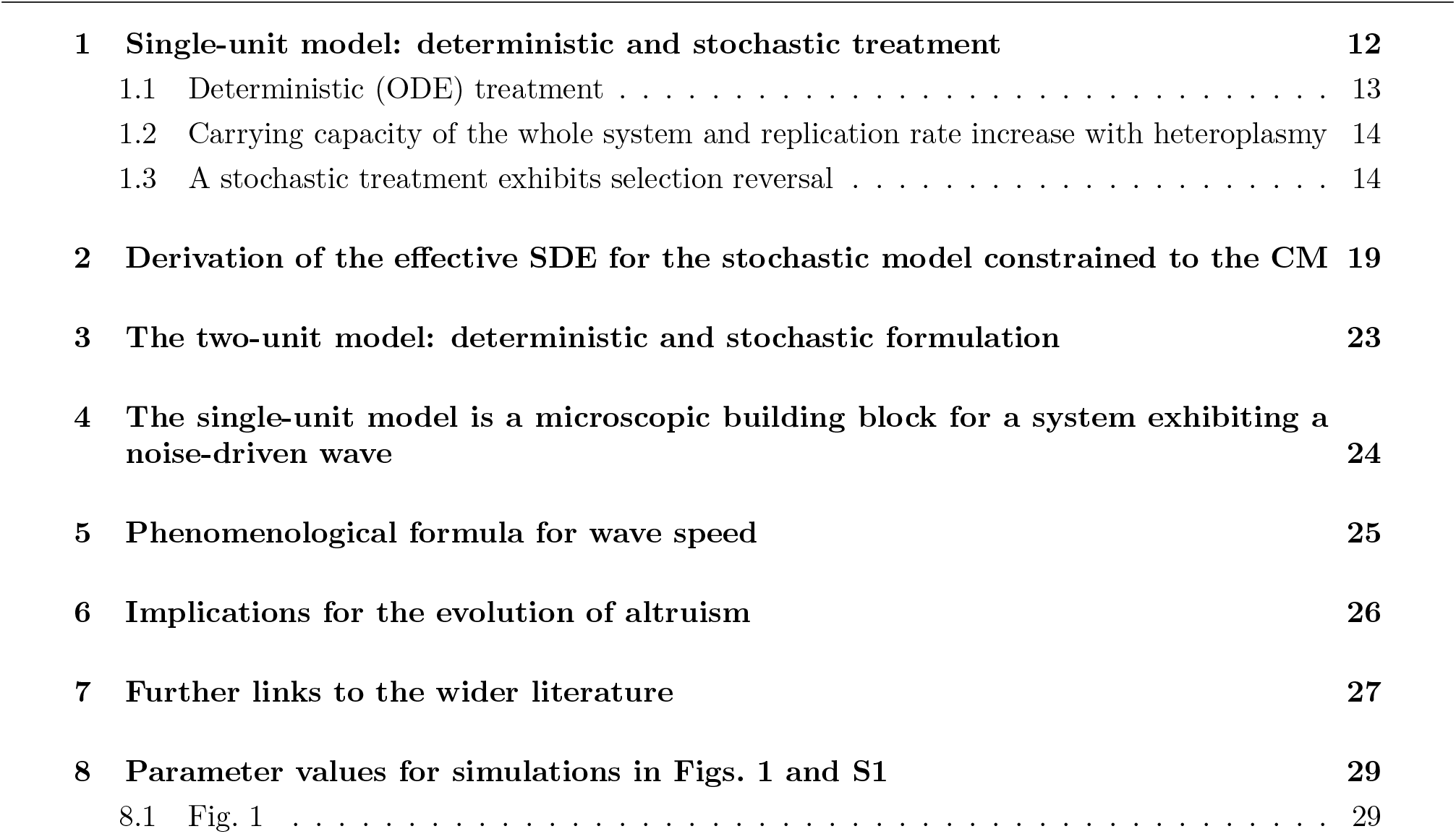
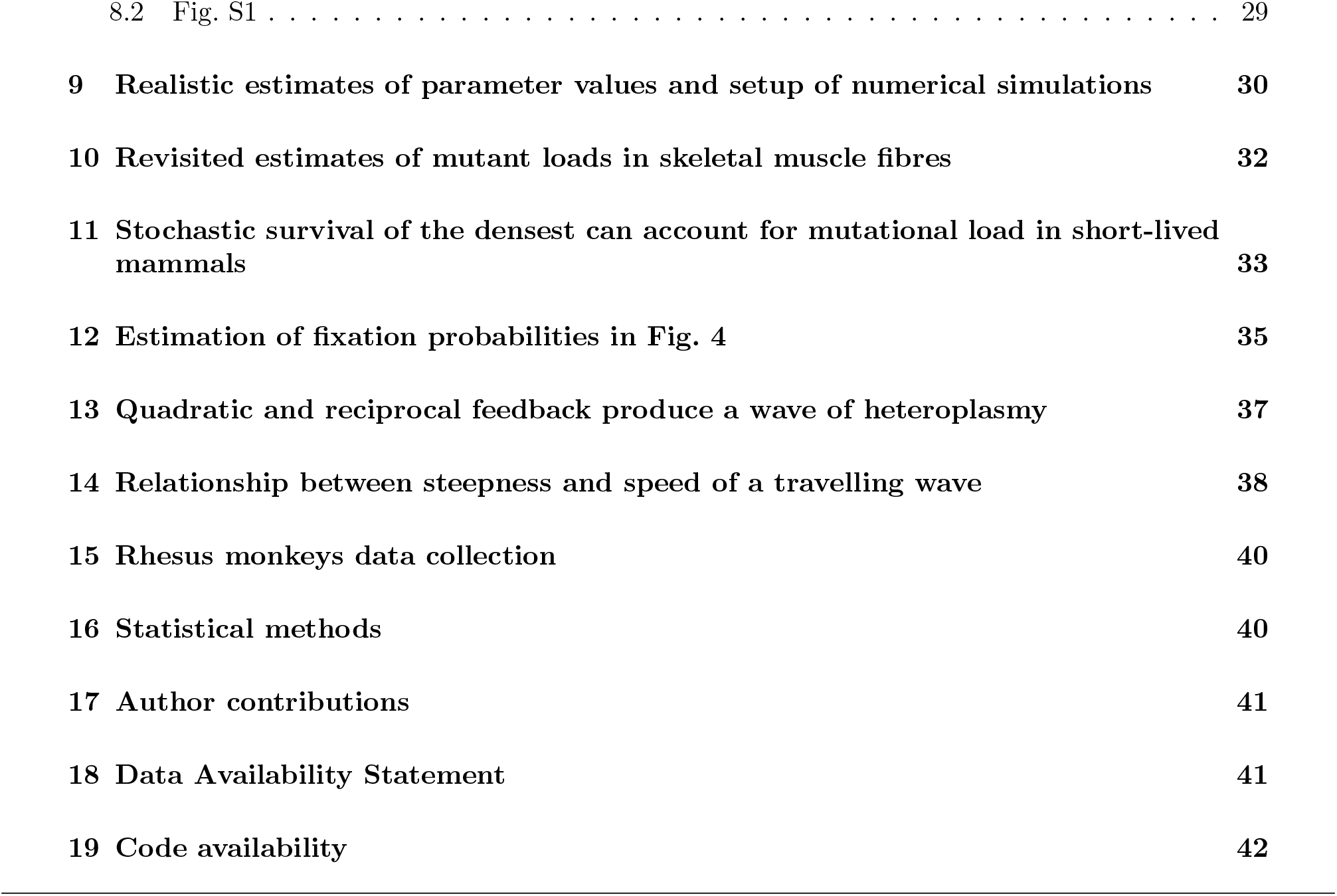
Supporting Information.

|  |  |  |
| --- | --- | --- |
| **1** | **Single-unit model: deterministic and stochastic treatment** | **12** |
| 1.1 | Deterministic (ODE) treatment . . . . . | 13 |
| 1.2 | Carrying capacity of the whole system and replication rate increase with heteroplasmy | 14 |
| 1.3 | A stochastic treatment exhibits selection reversal . . . . . | 14 |
| **2** | **Derivation of the effective SDE for the stochastic model constrained to the CM** | **19** |
| **3** | **The two-unit model: deterministic and stochastic formulation** | **23** |
| **4** | **The single-unit model is a microscopic building block for a system exhibiting a noise-driven wave** | **24** |
| **5** | **Phenomenological formula for wave speed** | **25** |
| **6** | **Implications for the evolution of altruism** | **26** |
| **7** | **Further links to the wider literature** | **27** |
| **8** | **Parameter values for simulations in Figs. 1 and S1** | **29** |
| 8.1 | Fig. 1 . . . . . | 29 |

|  |  |
| --- | --- |
| 8.2 Fig. S1 . . . . . | 29 |
| **9 Realistic estimates of parameter values and setup of numerical simulations** | **30** |
| **10 Revisited estimates of mutant loads in skeletal muscle fibres** | **32** |
| **11 Stochastic survival of the densest can account for mutational load in short-lived mammals** | **33** |
| **12 Estimation of fixation probabilities in Fig. 4** | **35** |
| **13 Quadratic and reciprocal feedback produce a wave of heteroplasmy** | **37** |
| **14 Relationship between steepness and speed of a travelling wave** | **38** |
| **15 Rhesus monkeys data collection** | **40** |
| **16 Statistical methods** | **40** |
| **17 Author contributions** | **41** |
| **18 Data Availability Statement** | **41** |
| **19 Code availability** | **42** |

## Introduction

In this Supporting information we provide arguments for the statements contained in the main text, derive the mathematical results underpinning this work, report further numerical results (detailing the setup of simulations) and provide estimates of the parameter values used for simulations shown in the main text.

In SI.1 and SI.2 we formulate the single-unit system and detail the stochastic dimensionality reduction procedure that leads to an effective single-SDE description of its dynamics. Understanding the single-system dynamics is the basis to account for stochastic survival of the densest (SSD), observed in stochastic spatially extended systems, such as the two-unit system formulated in SI.3. Further, in SI.4 we show mathematically that the single-unit system is the building block for a continuous model exhibiting a travelling-wave of mutants. In SI.5 we detail how we obtained the phenomenological wave speed formula Eq. (2) for our microscopic spatially extended model.

The following two sections position our work in the wider evolutionary biology literature. In SI.6 we define the contribution of this work to the debate on the evolution of altruism, while in SI.7 we differentiate SSD from density-dependent selection.

SI.8 details the parameter values used for simulations relating to some of the panels of Figs. 1 and S1, plots that do not refer to actual biological systems and are meant to illustrate SSD. In SI.9, instead, we provide estimates of the model parameters, derived from independent experimental studies.

One of the main contributions of this work is to provide a mechanistic accounts of the expansion of mitochondrial deletions in skeletal muscle fibres that can explain the observed mutant loads without requiring excessively high *de novo* mutation rates. In SI.10 we provide revised estimates of the mutant loads from experimental studies based on skeletal muscle fibres biopsies. In SI.11 we provide a mathematical argument that SSD can explain the observed loads with drastically lower mutation rates than previous models.

In SI.13 we demonstrate that the specific form of Eq. 1, with the replication rate *λ* depending linearly on copy numbers, is not necessary for SSD. The increase in mean heteroplasmy is observed with other replication rates that encode the higher density of mutants, provided that noise and spatial structure (with diffusion) are also present.

In SI.14 we provide computational evidence that, for waves driven by SSD, the steeper the wave-front, the slower the speed, a relationship that is also present in waves driven by a replicative advantage.

Finally, in SI.15 we explain how we obtained the data points plotted in Fig. 2B. and in SI.16 we define error bars in all our plots and give details on the statistical models and tests used.

## 1 Single-unit model: deterministic and stochastic treatment

*We formulate the single-unit model, in its deterministic (Section 1.1) and stochastic version (Section 1.3), and explicit the dependence of the system’s carrying capacity on mutant fraction (Section 1.2)*

The fundamental unit of our model is the single unit hosting a well-mixed population evolving according to linear feedback control. This constitutes the building block of our description of the skeletal muscle fibre. Our physical model of muscle fibres is a chain of these units.

In our model, there are two species: wildtypes *w* and and mutants (deletions) mitochondrial DNA molecules *m*.

Every molecule is degraded at rate *μ* and replicates at rate *λ*(*w, m*) given by

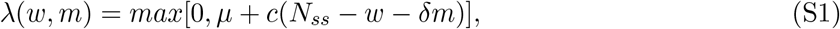

i.e. the larger between 0 and *μ + c*(*N_ss_* − *w* − *δm*). However, for biologically relevant values of the parameters, *μ + c*(*N_ss_* − *w* − *δm*) is practically always positive, as we noticed in our simulations. Therefore we can also use the simpler form in Eq. (1). Because of the linear relationship between the replication rate *λ* and the copy number we name the replication rate *linear feedback control*.

The meaning of the parameters *N_ss_* and *δ* are explained in the main text, where we also mentioned that *c* > 0 modulates the strength of the control. This means that the larger *c*, the more strongly the system is penalised to be away from steady state when *w* + *δm* ≠ *N_ss_*, i.e. when the terms in parentheses on the right-hand-side (RHS) is ≠ 0 and hence *λ* ≠ *μ* In other words, a larger value of *c* > 0 will cause a stronger push toward steady state.

In the following we study a deterministic and stochastic version of the model defined by the above rates.

### 1.1 Deterministic (ODE) treatment

In a deterministic setting, the above rates give rise to the following system of ODEs:

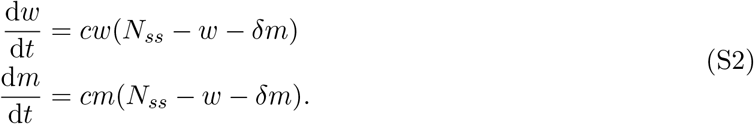

The analysis of this deterministic dynamical system is useful to understand the stochastic version of the model. We limit our analysis to *w* > 0, *m* > 0 given the meaning of the variables as copy numbers. We start by noticing the existence of the trivial, unstable fixed point (0, 0). We then notice a set of attractive fixed points which form the straight line of equation *w* + *δm* = *N_ss_* in the (*m, w*) plane. We refer to this as the central manifold (CM) or steady state (*μ* = *λ*) line. An example of CM, for a system with for a system with *N_ss_* = 500 and *δ* = 0.5, is plotted in Fig. S1A. The CM includes the configurations in which there is only a species, namely the wildtype fixed point (*N_ss_*, 0) and the mutant fixed point (0, *N_ss_*/*δ*). The *carrying capacity* of a species is the population size of this species at its fixed point, at which the species exists in isolation. We see that in our model the carrying capacities are *N_ss_* for wildtypes and *N_ss_*/*δ* for mutants. When 0 < *δ* < 1 – that is always the case in this work – the mutant carrying capacity is larger than wildtype. Another interpretation of the condition 0 < *δ* < 1 is that mutants are less tightly controlled or sensed by the system, as a change in the number of mutants entails a smaller variation in *λ*(*w, m*) than an equal change in the number of wildtypes. Hence, in our model mutants are the *densest* and *less tightly controlled* species. We define the *heteroplasmy h* as the proportion of mutants:

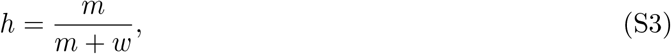

a conserved quantity of the dynamical system, namely

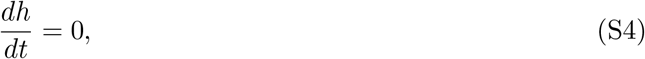

which can be verified by direct calculation. Eq. (S4) accounts for the constant heteroplasmy in the single-unit neutral deterministic model, illustrated in Fig. S1B (black line). Eq. (S4) is equivalent to the ratio *m/w* being constant, meaning that any line passing through the origin is a constant-heteroplasmy line. Hence, a way to summarise the behaviour of the system is:

- If the system is in steady state, i.e. if *w* + *δm* = *N_ss_*, the dynamics stop.
- If the system is not in steady state, it will move toward the CM along a line connecting the initial condition to the origin.

The same behaviour can be recovered from the full solution of the system, which is

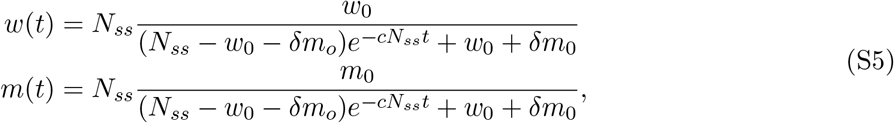

where *w*_0_ = *w*(0) and *m*_0_ = *m*(0) are the initial conditions. From Eq. (S5) it follows that

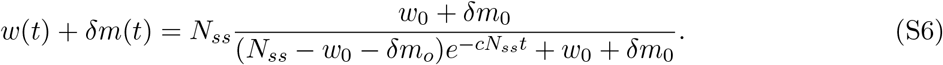

For *t* → ∞, as 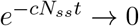, *w*(*t*) + *δm*(*t*) → *N_ss_*. This shows that the parameter *c* is connected to how fast the state of the system decays to the CM, justifying its interpretation of *c* as control strength.

In summary, in the deterministic single-unit model the system evolves toward the CM of equation *w* + *δm* = *N_ss_* with the condition *ḣ* = 0. We remark on the difference in carrying capacity of the two species, being modulated by the parameter *δ*.

### 1.2 Carrying capacity of the whole system and replication rate increase with heteroplasmy

The global carrying capacity *K* of the system is the value of the total population *n* = *w* + *m* at steady state. *K* can be expressed as a function of *h*. Since *m = hn* and *w* = (1 − *h*)*n* the steady state condition *w* + *δm* = *N_ss_* becomes

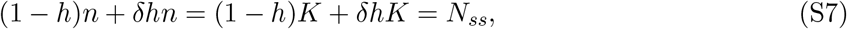

where the first equality holds because we have defined *K* as the value of *n* at steady state. This leads to

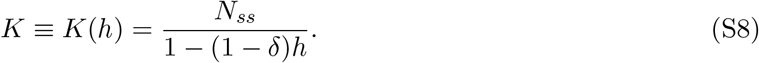

For *δ* < 1, *K*(*h*) is an increasing function of *h*: the higher the proportion of mutants, the larger the population that the system can sustain. When *h* = 1, i.e. mutants are in isolation, the mutant carrying capacity 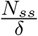 is recovered. One way to interpret this is that mutants are using the resources of the system in a more economical way. The economical use of a limited resource is considered one of the earliest and simplest forms of altruism ^33,56^, that brings benefits to all the other individuals in the system regardless of their identity.

Another way of seeing the benefit that mutants bring to all the other molecules in the system is in terms of enhanced replication rate. By expressing *w, m* in terms of *h, n* as above, Eq. (S1) can be written as

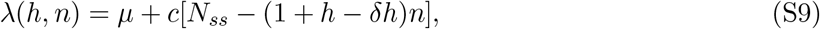

that shows that, for a given population size *n*, the common replication rate is an increasing function of *h* when *δ* < 1. Notice that here, differently from Eq. (S8), *n* ≠ *K* since we are not assuming that the system is in the steady state.

In the study of the expansion of mitochondrial deletions, mutants are assigned a higher degradation rate, to model the higher mitophagy rate to which they are subject. The mitochondrial deletions of our model, for which *δ* < 1, can hence be seen in a very general sense as agents that benefit others paying a cost, in the form of a higher degradation rate. This aligns with the definition of biological altruism ^56,57^, hence the mutants of our model can be considered an altruistic species.

### 1.3 A stochastic treatment exhibits selection reversal

In the stochastic formulation of the model, the *per capita* degradation and replication rates *μ* and *λ* are interpreted as instantaneous probability (in an infinitesimal time interval) that each molecule is degraded or replicates. In this setting, *w, m* and *h* are random variables, and we ask questions about their probability distributions and their moments. Stochastic population dynamics models can be formalised as chemical reactions networks, consisting, in the terminology of chemical systems, of a set of reactants, reactions and products. Consider a general chemical system consisting of *N* distinct chemical species (*X_i_*) interacting via *R* chemical reactions, where the *j*^th^ reaction is of the form

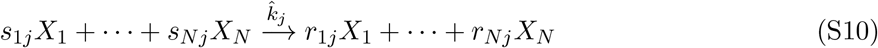

where *s_ij_* and *r_ij_* are stoichiometric coefficients. We define 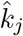 as the per molecule rate for the *jth* reaction. A chemical reaction network can be described by the associated *N* × *R* stoichiometry matrix *S_ij_ = r_ij_* − *s_ij_* and the set of rates 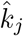. The reactants are the molecules of the two species. In our model the reactions are birth and death and the possible products are i) another molecule of the same species (birth), or ii) Ø for the degradation of a molecule (death reaction).

The chemical reaction network for the single-unit model of the first section of Results is

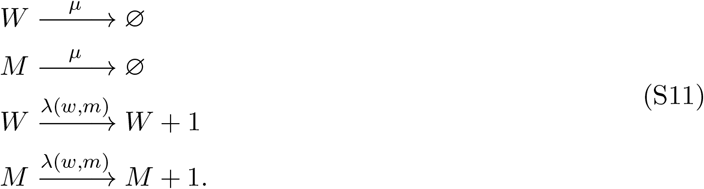

This is a network with *N* = 2 species and *R* = 4 reactions, with stoichiometry matrix given by

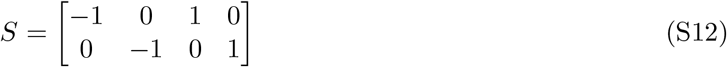

The global rates for wildtypes or mutants are found by multiplying the per molecule rates *μ, λ* by *w* or *m*. This is true only in the case *S_ij_* ≤ 1; more details for the general case can be found in Ref. [19]. These reactions and rates can be used to set up a Chemical Master Equation (CME), a system of coupled ODEs in *P* (*w, m, t*), the probability that the system is in the state (*w, m*) at time In principle, solving the CME would give the probability that the system is found in any state at any given time. However, given the nonlinearity of the global rates, the CME cannot be solved exactly. One can explore the behaviour of the stochastic models by simulating the CME, which can be done through Gillespie’s stochastic simulation algorithm^61,62^. This algorithm is exact, in that each Gillespie simulation represents an exact sample from *P* (*w, m, t*). The simulations allowed us to observe the increase in mean mutant copy number (*m*) for *δ* < 1 (Fig. S1B, red line). Intuitively, while fluctuating around the steady state line the system on average drifts toward regions of higher *m* (Fig. S1A).

It is worthwhile having an analytical account of this effect, in order to establish how the increase in mutant copy number depends on the parameters of the model. We have obtained an effective description of the system, in the form of an approximate stochastic differential equation (SDE) that shows an increase in the number of mutants for *δ* < 1. The steps are the following:

1. Applying the Kramers-Moyal expansion to the CME, obtaining a Fokker-Planck equation (a PDE for the probability distribution *P* (*w, m, t*)).
2. Converting the Fokker-Plack Equation into a system of two coupled SDEs in the variables *w*(*t*) and *m*(*t*).
3. Applying a stochastic dimensionality reduction procedure, that exploits the fact that the system fluctuates around a central manifold (CM), to get a single SDE for *m*(*t*) that shows a positive drift for *δ* < 1.

The first two steps are standard ^63,64^. From the chemical reaction network in Eq. (S11), we obtain the system of SDEs:

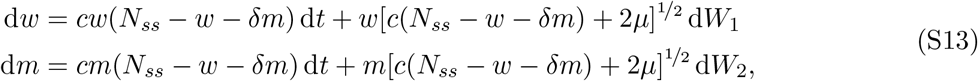

where d*W*_1_ and d*W*_2_ are two i.i.d. Wiener increments, i.e. Gaussian noise with zero mean and variance *dt* (see also Eq. S31). The third steps relies on a recently developed technique^30,65,66^, which works specifically for systems fluctuating around a CM. In the next section SI.2 we detail each step, while here we only present and interpret the results. The final, effective SDE is

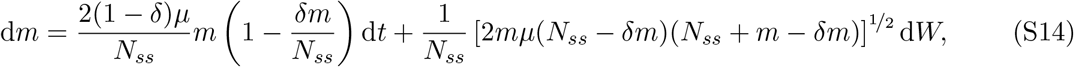

with *dW* a Wiener increment. The first term on the RHS is the drift, and it corresponds to a logistic growth with carrying capacity 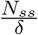 When the drift is positive, since 〈*dW*〉 = 0, copy number increases on average. The drift is positive for 0 < *δ* < 1, i.e. when mutants have a higher carrying capacity or density, and *δm* < *N_ss_*. Because of noise the final value of the mutant copy number will not be 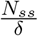.The dynamics stop either when *m* = 0 or 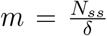, the only values for which *dm* = 0. This approximation relies on the system fluctuating closely around the CM, which is true for large values of *c* (strong control). The parameter *c* is not present in Eq. (S14) because this equation holds exactly for *c* → ∞. Indeed, when simulating Eq. (S14) using the Euler scheme, the agreement with the results of the exact Gillespie simulations is better for larger values of *c* (not shown). By applying Itô’s formula (see SI.2) it is possible to find the SDE for *h*, which is

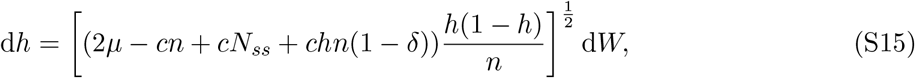

where *n* = *w* + *m*. We notice in this equation the absence of a drift term, meaning that

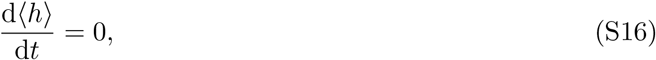

i.e. mean heteroplasmy is constant in the single-unit stochastic model, as stated in the main text. This is shown in Fig. S1C, that reports data from an ensemble of stochastic simulations (apart from the black line that refers to the ODE system).

A heuristic way to understand why the mean number of mutants 〈*m*〉 increases and mean hetero-plasmy 〈*h*〉 is constant is the following. The per capita rates in Eq. (S11) are linear, hence the system is a stochastic Lotka-Volterra model, for which each individual has the same chance of generating a lineage that will take over the whole system^67^. Hence, this probability must be 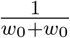 Since there are *m*_0_ mutants, the probability that a mutant will take over the whole population is 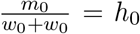 : the mutant fixation probability coincides with the initial fraction of mutants. At mutant fixation, the steady state mutant population is 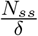 and the mutant fraction is (obviously) 1. Based on this, one can conclude that, for *t* → ∞, 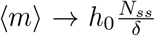 and 〈*h*〉 → *h*_0_ · 1 = *h*_0_. Taking the limit *t* → ∞ means that we wait long enough for fixation to happen and for the population to reach the steady state. Recall that this differs from the deterministic case, in which there is a locus of fixed points of equation *w* + *δm* = *N_ss_* (the CM) and the system equilibrates at the point of the CM for which heteroplasmy is *h*_0_. For 0 < *δ* < 1, 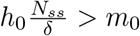 always, hence 〈*m*〉 will increase to reach its final value. Conversely, the final value of 〈*h*〉 coincides with the initial value *h*_0_, giving intuition as to why 〈*h*〉 is constant. It is also illuminating to consider the case in which *δ* ≈ 0, namely when the mutants are scarcely controlled. When a wildtype dies at steady state this increases the birth rate. Either a mutant or a wildtype will be the next birth, but if a mutant is born this leaves the elevated birth rate (caused by the degradation of the wildtype) almost unchanged, encouraging further possible mutant births. Interestingly, even for *δ* ≈ 0 and in the extreme, unrealistic case of infinite mutant density *δ* = 0, 〈*h*〉 stays constant. Indeed, Eq. (S16) and the above heuristic argument are valid for any value of *δ*.

Importantly, the stochastic system can exhibit an increase in 〈*m*〉 even if we introduce preferential elimination of mutants (Fig. S1B, green line). This can be done by increasing mutant degradation rate by *ϵ*, reflecting what we expect for defective mtDNA molecules. Indeed, under certain cases there is evidence for enhanced clearing of defective mutations ^68,69^, subject to higher degradation rates. Mathematically, this is described by replacing the second reaction in Eq. (S11) by

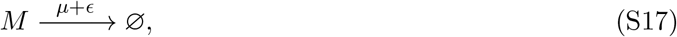

with *ϵ* > 0, leaving all other reactions unchanged. In the next section (Eq. (S34)) we will see that, through the techniques in Refs. [30, 65, 66], it is possible to derive an SDE for mutant copy number that is valid for *ϵ* ≪ *μ* :

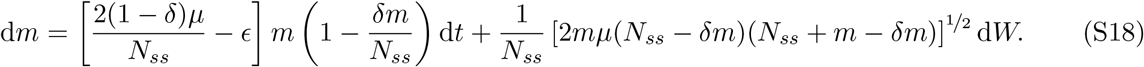

The drift in *m* is now positive for *δm* < *N_ss_* and 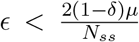. In this case, the combined effect of stochasticity and differences in carrying capacity (density) produces a stochastic reversal of the direction of deterministic selection ^30^. Indeed, in the deterministic case, for any *ϵ* > 0 and for any initial condition with non-zero *w*_0_ the system tends to the point (*N_ss_*, 0), the only stable fixed point, corresponding to wildtype fixation (mutant extinction). Conversely, in the stochastic case, there is a critical 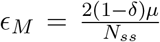 such that, for *ϵ* < *ϵ_M_*, (*m*) still increases, meaning that mutant fixation is more likely than wildtype. Inspection of the expression for *ϵ_M_*, interpretable as the largest amount of additional mitophagy rate that still allows mutant to expand, reveals the following interesting points:

- *ϵ_M_* increases with *μ*, meaning that the noisier the system, the higher the mitophagy rate that mutants can tolerate and still expand.
- For 0 < *δ* < 1, *ϵ_M_* is decreasing function of *δ*. The higher the difference in density, the higher the mitophagy rate that mutants can overcome.
- *ϵ_M_* is a decreasing function of *N_ss_*. In particular, *ϵ_M_* → 0 when *N_ss_* → ∞. This is common to stochastic effects, whose intensities decrease with the system size, vanishing in the deterministic limit for which system size tends to infinity.

## 2 Derivation of the effective SDE for the stochastic model constrained to the CM

*We derive an effective SDE in the single-unit system, that accounts for the increase in mean copy number even in the presence of a higher mutant degradation rate*

In this section we show that Eq. (S14) is an effective, approximate description for the system defined by the chemical reaction network in Eq. (S11) given the replication rate in Eq. (S1). We provide detail for each of the steps listed in SI.1. The approach used here is adapted from standard texts ^63,64^ and from Refs. [65, 66]. We strive to provide a self-contained treatment, hoping to make our derivation and the set of techniques accessible to a wider audience. Our exposition partly follows Ref. [19], where the same procedure is used for a related problem.

### 1- From the CME to a Fokker-Planck equation via Kramers-Moyal expansion

Let us denote **n** = (*w, m*), a vector specifying the system state. The CME can be written in compact form as

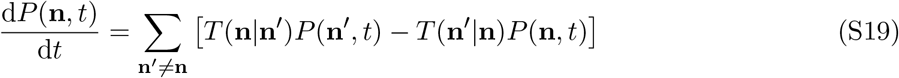

*T* (**n**|**n**′) is the transition rate from state **n**′ to **n**. Given the reaction network in Eq. (S11), the corresponding (global) transition rates are

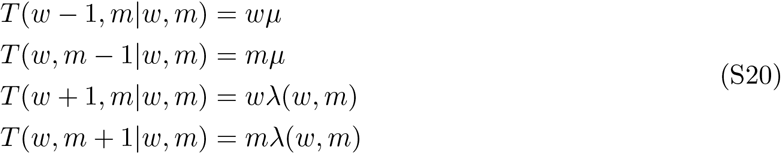

The first step is to write the CME in a continuous form. We replace copy numbers **n** by the abundances 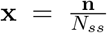 relative to the effective population size. The variable **x** can be considered continuous for large *N_ss_*. The CME becomes

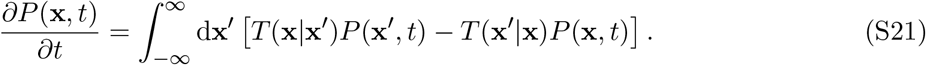

We now proceed by expanding the CME through the second-order multivariate Kramers-Moyal expansion ^63^, that can be written as

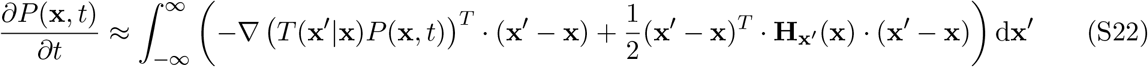

where **H_x′_** (**x**) is the Hessian matrix of *T* (**x**′|**x**)*P* (**x**), whose general expression is

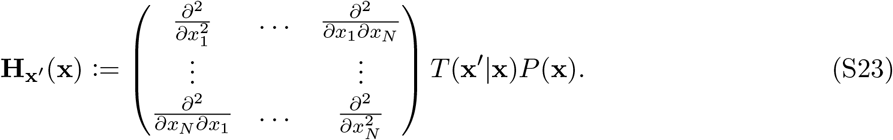

A transition **x** → **x**′ corresponds to some reaction *j* which moves the state from **x** to **x**′. Since we know how each reaction affects state **x** through the constant stoichiometry matrix *S_ij_*, and since the global rate of each reaction is independent from **x**′ itself (see Eq. (S20)), we may transition from a notation involving **x** and **x**′ into a notation involving **x** and the reaction *j* that affects **x**. We may therefore define *T_j_*(**x**) := *T* (**x**′|**x**), and let **H_x′_** (**x**) → **H***_j_*(**x**).

We now wish to re-write Eq. (S22) as a Fokker-Planck equation. Since the integral in Eq. (S22) is over **x**′, and for a given **x** every **x**′ corresponds to a reaction *j*, we may interpret the integral in Eq. (S22) as a sum over all reactions, i.e. 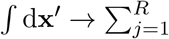. Hence, for the *j*^th^ reaction, [(**x**′ − **x**)]*i* = *S*_*ij*_. With these observations, we may write the first term in the integral in Eq. (S22) as

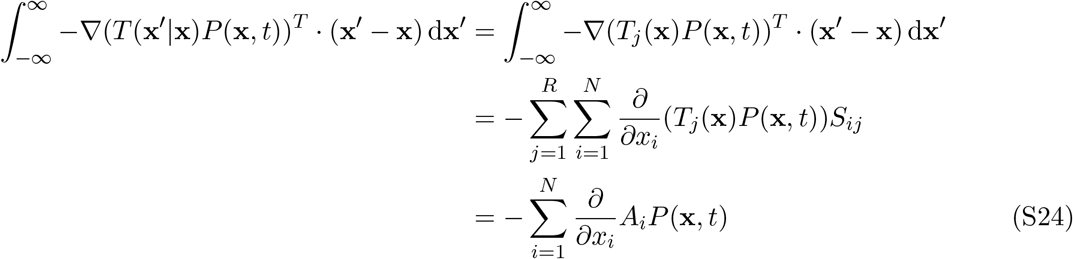

where

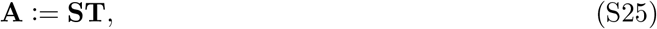

with **A** a vector of length *N*, *S* the *N* × *R* stoichiometry matrix and **T** the vector of transition rates, of length *R*. To re-write the second integral of Eq. (S22), we write an element of the Hessian **H***_j_* in Eq. (S23) as

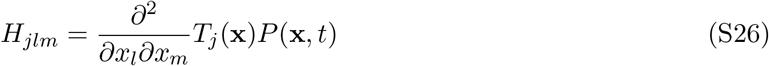

where *j* = 1, …, *R* and *l, m* = 1, …, *N*. Thus, we may write

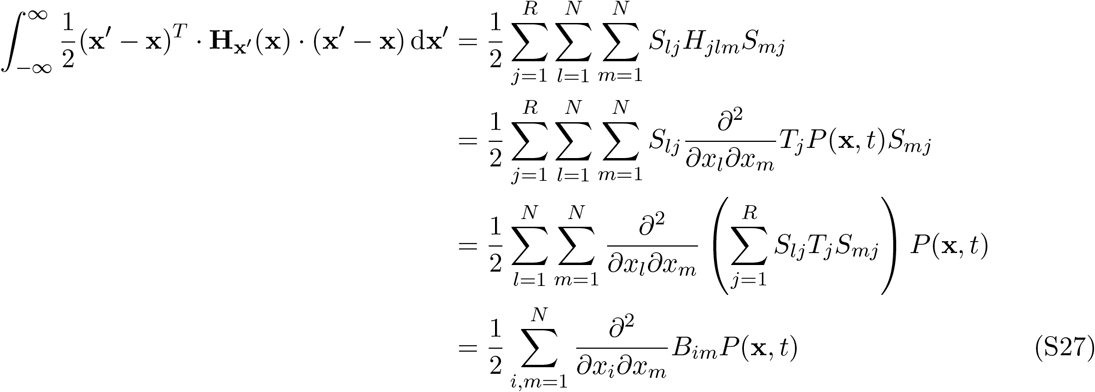

where

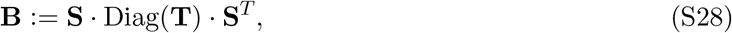

and **B** is an *N* × *N* matrix, and Diag(**T**) is a diagonal matrix whose main diagonal is the vector **T**. We may therefore re-write Eq. (S22) as a Fokker-Planck equation for the state vector **x** of the form

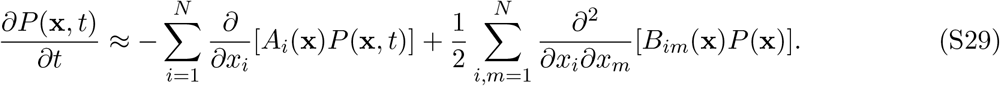

From a system of coupled ODEs for *P* (*w, m, t*) (the CME of Eq. (S19)) we have obtained a single PDE for *P* (**x**, *t*) approximating copy numbers as continuous variables.

### 2- From Fokker-Planck to a system of Itô SDEs

In general, the Fokker-Planck equation in Eq. (S29) is equivalent^64^ to the following Itô stochastic differential equation (SDE)

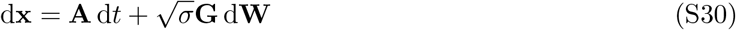

where **GG***^T^* ≡ **B** (where **G** is an *N* × *R* matrix) and d**W** is a vector of length *R* of independent Wiener increments, that satisfy

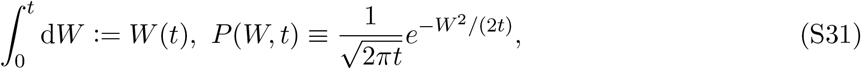

and *σ* is a constant that controls the strength of the noise. For our chemical reaction network Eq. (S11), the corresponding system of Itô SDEs is given in Eq. (S13).

### 3- From a system of Itô SDEs to a single effective SDE for a system forced onto the central manifold

What follows is an adaptation from ^65,66^, to which we refer for a proof and for interpretation of the functions introduced. We start from Eq. (S30). The procedure can be applied when the system admits a manifold on which **A** = 0, the central manifold (CM). We though it would be an helpful complement to Refs. [65, 66] to give an explicit sequence of the steps involved for the specific system at hand.

- Identify the CM.
- Find the Jacobian of **A**, namely *J* such that

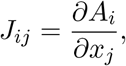

and evaluate it on the CM, obtaining *J_CM_*.
- Find the eigenvalues *λ*_1_, · · ·, *λ_N_* of *J_CM_*. The CM is the kernel of *J_CM_*, hence the multiplicity of the zero eigenvalue is the dimensionality of the CM.
- Compute the decomposition of *J_CM_*, writing

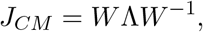

where *W* = (**w_1_**, · · ·, **w_N_**) is the matrix of eigenvectors with those corresponding to the zero eigenvalues written first.
- Compute 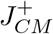, defined as

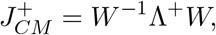

where Λ^+^ is the diagonal matrix with diagonal elements 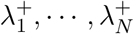 defined as

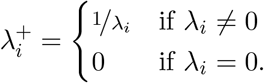
- For each element of **A**, compute the Hessian *H_i_* as

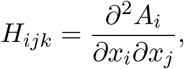

and evaluate it on the CM, obtaining *H_i_CM*.
- Compute the matrix 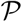 as

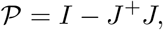

where *I* is the identity matrix, and the matrices *Q_i_* as

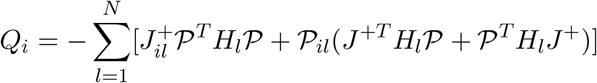
- Calculate the vector **g**(**x**), whose elements are

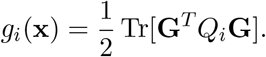
- Finally, effective SDEs for the variable **z**, namely the variable **x** constrained to the CM, are

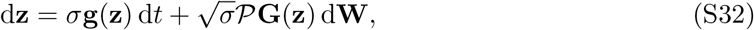

where the equations are now *uncoupled* because all the functions are evaluated on the CM.

This procedure allows one to obtain Eq. (S14) as an effective description of the neutral stochastic model defined in Eq. (S11). We provide a Mathematica notebook (see Supplementary File 1) to obtain Eq. (S30) from the system of SDEs in Eq. (S13). This technique is extended in Ref. [65] to the more general case in which the stochastic system is equivalent to a system of Itô SDEs of the form

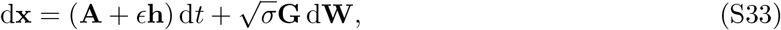

where **h** ≡ **h**(**x**) is another vector-valued function of **x** and *ϵ* ≪ *σ* is a small parameter, such that for

*ϵ* = 0 the system admits a CM. In this case, Eq. (S32) becomes

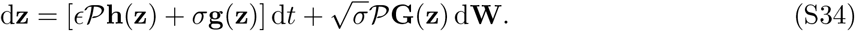

The stochastic system in which mutants are subject to preferential elimination *ϵ* (Eq. (S17)) corresponds to the case **h**(**x**) = (0, *x*_2_) (recall that *x*_2_ = *m/N_ss_*). Inserting this into Eq. (S34), one obtains Eq. (S18), that describes the noise and density-induced selection reversal.

### Change of variable through Itô’s formula to obtain an SDE for heteroplasmy

Itô’s formula states that, for an arbitrary function *y*(**x**, *t*) where **x** satisfies Eq. (S30), we may write the following SDE:

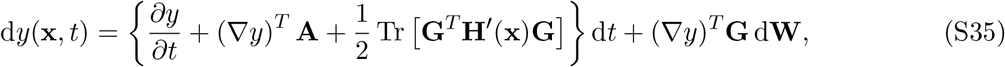

where **H**′(**x**) is the Hessian matrix of *y*(**x**, *t*) with respect to *x* (see Eq. (S23), where *T* (**x**′ | **x**)*P* (**x**) should be replaced with *y*(**x**, *t*)). Applying this rule to Eq. (S14) for

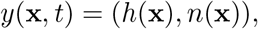

where *h* is the heteroplasmy and *n* is the total copy number, one obtains Eq. (S15)

## 3 The two-unit model: deterministic and stochastic formulation

*We formulate the deterministic and stochastic two-unit model, the latter being the simplest system exhibiting stochastic survival of the densest*

The deterministic two-unit model is formalised via the ODE system

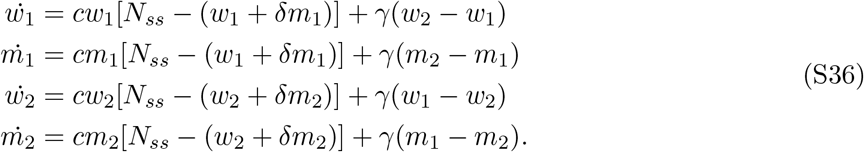

The addition of the second term on the RHS, proportional to the constant hopping rate *γ* accounts for diffusion of the mtDNA molecules between neighbouring units. The first term on the RHS of each equation means that each cell seeks to maintain its own separate target population, i.e. copy number control is local. This system does not admit an analytical solution. However, it has a CM given by *w*_1_ + *δm*_1_ = *w*_2_ + *δm*_2_ = *N_ss_*, *w*_1_ = *w*_2_ (and consequently *m*_1_ = *m*_2_). It is easy to see that if these conditions are satisfied, all the time derivatives (the left-hand-sides) are zero and the dynamics stop, i.e. steady state is reached. Afterwards, heteroplasmy stays constant, as shown in Fig. S1D (black line). A similar system of ODEs can be used to describe chains of units of arbitrary length (see SI.5).

The stochastic formulation of the model is obtained by replicating the reaction network in Eq. (S11) for *w*_2_, *m*_2_ and adding the reactions accounting for diffusion of the mtDNA molecules. The death rate is still constant and common, and each unit has its own independently controlled replication rate given by *λ_i_* = *μ* + *c*(*N_ss_* − *w_i_* − *δm_i_*), for *i* = 1, 2. The full chemical reaction network is

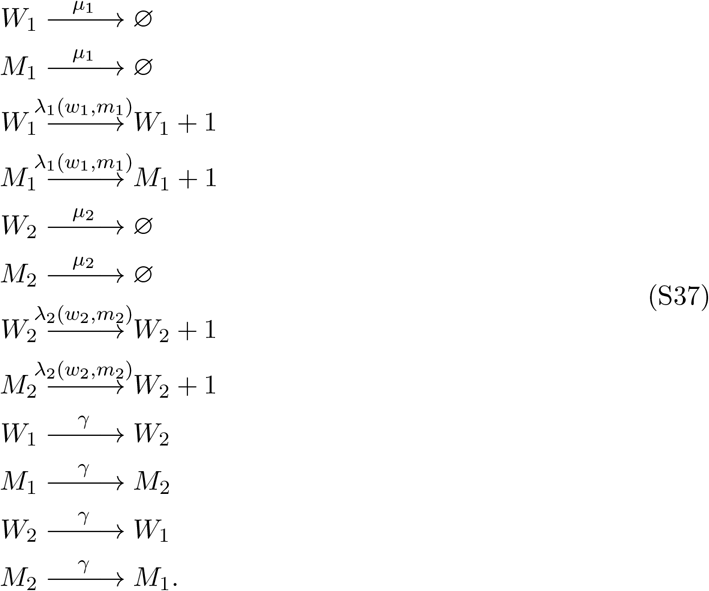

The dimensionality reduction technique from^65^, used to obtain Eq. (S18) is not viable in this case to obtain an SDE showing the increase in 〈*h*_1_〉 (or 〈*h*_2_〉) when 0 < *δ* < 1. The technique works by projecting the system onto the CM, where the two units are identical; however, the increase in mean heteroplasmy is driven by fluctuations that move one of the units away from the CM condition, after which the system relaxes to steady state with an average increase in *h*. The approximation involved in the technique is too crude to capture the effect. However, stochastic simulations consistently show the increase in 〈*h*〉 in chains of arbitrary length when 0 < *δ* < 1. This effect, conveniently highlighted in the two-unit system (see Fig. S1D, red line), is key to the appearance of a travelling wave of mutants in a chain of units. Crucially, the increase in mean heteroplasmy is observed even if mutants are subject to a slightly higher degradation rate (Fig. S1D, green line). This leads to a travelling wave of mutants, despite their preferential elimination, in a chain of units.

## 4 The single-unit model is a microscopic building block for a system exhibiting a noise-driven wave

*We show that a previous continuous model producing a noise-driven wave is a particular case of our microscopic mechanistic model*

In Ref. [33], a stochastic continuous-space model is presented, in the form of partial differential equations, describing a noise-driven wave of mutants, possibly subject to higher death rates than wildtypes. The paper is an extension of the classical Fisher-Kolmogorov PDE to a stochastic setting. In this section, we show how that our microscopically interpretable bottom-up model, based on the simple Lotka-Volterra dynamics, gives rise to the model in Ref. [33] for *δ* → 1 and in a continuous-space limit.

Let us reconsider the deterministic system of Eq. (S2) and change variable to (*n, h*), where *n* = *w* + *m* is the total population. We already observed that 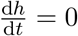. For *n* we have

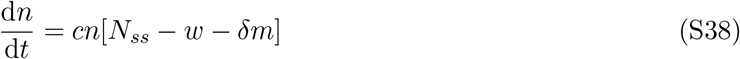

Let’s write *δ* = 1 − *α* :

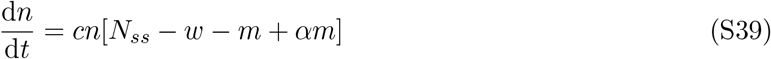

The constancy of *h* allows us to write *m* = *hn*, hence

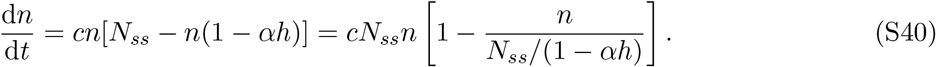

Considering the case *α* → 0, equivalent to *δ* → 1, we can write

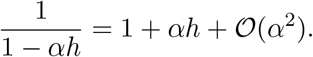

Neglecting 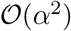 terms we obtain

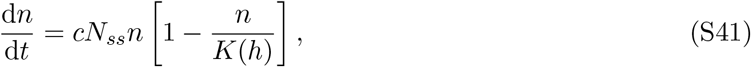

with *K*(*h*) = *N_ss_*(1 + *αh*). The total copy number follows a logistic growth with a carrying capacity *K*(*h*) that is a linearly increasing function of heteroplasmy *h*. In order to model a spatially-extended system as a muscle fibre, we give spatial dependence to copy number, i.e. *n* ≡ *n*(*x*). The hopping of mtDNA molecules between neighbouring units of the muscle fibres is described in this case as a diffusion term, proportional to the second spatial derivative of 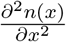 In order to model a muscle fibre, we can therefore write

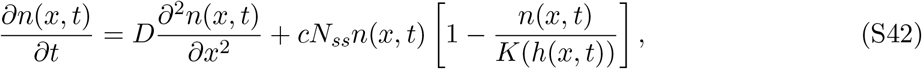

where we have explicitly written the time-dependence. *D* is the diffusion coefficient, for which *D = γL*^2^, where *L* is the inter-unit spacing.

Eq. (S42), derived from the special case of the simple single-unit Lotka-Volterra model, is the starting point of the work in Ref. [33]. By introducing noise from Wright-Fisher sampling and taking the low-diffusion limit *D* → 0, a wave equation for *h* can be derived which holds also in the case of slow preferential elimination of mutants. The limit *δ* → 1, that we have taken here to derive Eq. (S42) from Eq. (S2), is needed also in Ref. [33] in order to derive the wave equation. Hence our model, for some limiting values of the parameters *D* and *δ*, leads to a wave-equation for *h*. We stress the remarkable fact that from one of the simplest models of population genetics, the Lotka-Volterra, it is possible to derive an equation for a travelling wave of heteroplasmy driven by stochastic survival of the densest, albeit in some limit.

## 5 Phenomenological formula for wave speed

*We describe how Eq. (S43) was obtained and compare it to the wavespeed of the classical Fisher-Kolmogorov wave*

In the previous section we have shown how our model, in the limits *γ* → 0 and *δ* → 1, leads to an analytical description of a noise-driven wave. Moreover, through numerical simulations we have shown that the wave-like expansion is exhibited by our model without having to assume the above limits. We have performed simulations of a 201-unit chain with 110 combination of the parameter values, with *δ* ∈ [0.1, 1], *N_ss_* ∈ [2, 35], *γ* ∈ [0, 0.25], *μ* ∈ [0.01, 0.15]. For each parameter configuration we have observed a wave-like expansion, except, as expected, when *δ* = 1 or *γ* = 0. We have calculated the wave speed plotting the wavefront at different times and measuring the distance covered, measuring the degree of shift between the two wavefronts. In some cases, the overlap could not be total since the wavefront’s steepness changed slightly over time: in those cases we have looked for an overlap of the midpoint of the curves (i.e. the point for which 〈*h*〉 = 0.5). In all cases, it has been verified that the speed is constant, by measuring it for several different time intervals. We have found that *v* can be predicted with excellent accuracy through the formula

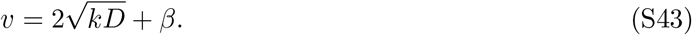

Apart from the intercept, Eq. (S43) is analogous to the formula for the wave speed of FK waves, with *k* being an effective reaction rate induced by stochastic survival of the densest, given by

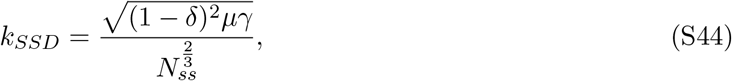

and *β* = 0.207 ± 0.007 is calculated via MLE (uncertainty is standard deviation). In Fig. S2 we show the fit, for which *R*^2^ = 0.99, and we observe residuals without any clear pattern. However, the presence of an intercept *β* ≠ 0 means that Eq. (S43) is not accurate for *x* → 0, for which *v* = 0. In Fig. 2F we compare the probability distribution of the wave speed predicted through Eq. (S43), obtained through Monte-Carlo sampling on the basis of the estimated distributions for the parameters of our model, given in SI.9.

Finally, we have verified that in a deterministic model deployed on a chain, i.e. Eq. (S36) applied to a larger number of units, the high-heteroplasmy front diffuses away even if *δ* < 1 (Fig. S4A), which is consistent with the statement that the wave-like expansion is noise-driven.

## 6 Implications for the evolution of altruism

*We connect our work to the debate on the evolution of biological altruism*

An altruistic trait has been defined as one that benefits others while costing its carriers ^56,58^. In SI.1.2, specifically in Eq. (S8), we show that the global carrying capacity of the system – the total population it can sustain – increases with the proportion of mutants. In this sense, mutants are a density-increasing species. In Eq. (S9) we show that, equivalently, the replication rate of both wildtypes and mutants is an increasing function of the proportion of mutants. If we assign a higher degradation rate to mutants, as we do in order to model a notional higher degradation rate for mitochondrial deletions, then our model can be linked to the specific definition of altruism in Refs. [56, 58]. Indeed, the increase in carrying capacity and replication rate can be seen as a benefit that mutants bring to the whole population, including wildtypes, at the cost of a higher degradation rate.

An increase in the system’s carrying capacity is present also in mathematical models of public good production ^30,70^ and cooperative use of limited resources ^56^. These two behaviours are observed in microbial communities and have been described as early examples of altruism in the history of life ^33,71^.

The two most prominent accounts for the evolution of altruism are kin selection and group selection ^72^. The former approach argues that altruism directed towards genetically related individuals can increase the overall success of the altruistic gene, which is likely to be shared by relatives of the carriers ^73^. In group selection, altruism spreads through advantages conferred to the group; for instance, a group whose members are ready to sacrifice themselves for other members will likely prevail over other groups ^74^. These approaches seek a *deterministic* advantage for altruistic traits. In our model, the spread of density-increasing mutants is *driven by noise* and has no deterministic counterpart. We now highlight our contributions in this area, referring to SI.7 for further links to the wider evolutionary biology literature.

A first, important merit of our model consists in describing the spread of the density-increasing trait in a spatially extended system using a fundamental and widely-used model of population genetics, the Lotka-Volterra model, which uses linear per molecule transition rates. The Moran model is another fundamental model of stochastic population genetics, dealing with the less general case of finite fixed-size populations. Previously, the study in Ref. [57] has shown that in a modified Moran Model with nonlinear rates, where the assumption of fixed population size is relaxed, density-increasing mutants (termed altruists by the author) can have a higher fixation probability despite a higher death rate. Analogous results are presented in Ref. [75], which is based on a modification of the Wright-Fisher model, retaining the feature of discrete, non-overlapping populations. Our continuous-time model uses simpler (linear) rates, and the relevance of the Lotka-Volterra model, widely used in population genetics, indicates that SSD might be a widespread effect.

The increase in mean mutant copy number in the single-unit system in a neutral model was highlighted 20 years ago in the field of mitochondrial biology, in the context of the relaxed replication model formulated in Refs. [11, 15, 16, 29] and further analysed in Refs. [29, 76]. Models yielding an increase in mean *copy number* of a denser mutant species have found that space can amplify the effect of randomness^30,65,66^, in the sense that the denser mutants can tolerate a higher selective elimination in a spatially structured system. Crucially, our work explicitly shows that spatial structure can lead to a qualitatively different result, namely an increase in the *proportion* of the density-increasing mutants, that in turn leads to a travelling wave invading the system.

Another contribution of our study is the use of a physically motivated and biologically interpretable microscopic mechanism. The study in Ref. [33] presents a phenomenological, macroscopic model that exhibits a noise-driven wave of (what the author calls) altruists. We show in the SI.4 that the model in Ref. [33] is a special case of ours, in the limit of continuous space. We have further been able to obtain a formula for the wave speed covering a range of biologically and clinically relevant regimes not covered by other studies, namely without taking *δ* → 1 and without imposing the quasi-static limit, i.e. very slow diffusion (see SI.5).

Finally, but perhaps most importantly, we have identified the mtDNA populations in skeletal muscles as a candidate system that shows a noise-driven travelling wave of a preferentially eliminated but density-increasing species. Muscle ageing could therefore be the first example of a new class of evolutionary phenomena best described as SSD.

## 7 Further links to the wider literature

*We differentiate our contribution from previous work on density-dependent selection and the impact of spatial structure on the evolution of altruism*

Density-dependent selection refers to situations in which the fitnesses of the different species within a population depend differently on population density (or size) ^77^. The theory of *r/K* selection ^78,79^, one of the most influential ideas in evolutionary biology ^80–82^, examines how selection shapes the strategies of species in the presence (or in absence) of density effects. The study of density-dependent selection has attracted interest across five decades ^78,83–85^, including in systems where stochasticity is important^77,86–91^. In stochastic survival of the densest, wildtypes and mutants have a common per capita replication rate. Per capita degradation rates are constant, and the mutant degradation rate can be higher than the wildtype by a constant additive factor. The mutant and wildtype rates therefore depend in the same way on copy numbers, and hence our model is not an example of density-dependent selection. Frequency-dependent selection ^92^ is distinct, but related, to density-dependent selection. Numerous definitions of frequency dependent-selection have been used over the decades, a classic one being that the relative fitness of a species varies with the relative frequencies of other species ^93^. Our model satisfies a strong mathematical definition of frequency-*independent* selection, based on the per capita growth rate (PCGR) of the species, the difference between per capita birth and death rates. The PCGR can be defined as a vector-valued function of copy number variables, with an output for each species. Frequency-independent selection is assured when the Jacobian of the PCGR is independent of copy numbers ^94^. This is clear for our linear feedback model (and all generalised Lotka-Volterra models), since the replication rate in Eq. (1) is linear in *w* and *m*. Therefore, stochastic survival of the densest is not a case of frequency-dependent selection.

In the context of a single-species population, the Allee effect can be described as a positive dependence of the PCGR on the density or size of the population for some interval of the values of population size ^95^. This contrasts with theories in which the PCGR always decreases with population size because of competition for resources or space among members of the same species (intra-specific competition), as in logistic growth. When the dynamics of a population exhibits the Allee effect, it means that there is some other mechanism at play that counterbalances the general limiting effect of competition on the PCGR. An example is that of plants that reproduce through pollination, whose rate is increased by the density of plants in the environments. The Allee effect is connected to altruism, in that the efficiency of some fitness-increasing behaviours, e.g. defence and feeding, is enhanced by a large population size when the associated tasks are carried out cooperatively ^96^. The difference between stochastic survival of the densest and the Allee effect is two-fold. The Allee effect refers to the possibility that PCGR in a population can be an increasing function of population size (for some interval) despite the increase in intra-specific competition that generally comes with higher density. Our work instead models inter-specific (between species) competitions, showing that the species that is *denser in isolation* can prevail despite a higher degradation rate and no replicative advantage. Moreover, the PCGR of our neutral model is proportional to *N_ss_* − *w* − *δm* for both species; in the case of a higher degradation rate for mutants, their PCGR is proportional to *N_ss_* − *w* − *δm*. In both cases we have linearly decreasing functions of the population size, with no positive dependence on it.

Previous work has explored how the spatial structure of a population can favour ^97,98^ or hinder ^99^ the evolution of altruistic behaviour. These studies refer to game theoretical models that can be recast in terms of population dynamics models, due to the correspondence between the replicator equation and the generalised Lotka-Volterra model, in their deterministic ^100^ and stochastic ^101^ versions. Similar results have been found applying agent-based models to cell populations ^56,102^. In these studies, the *continuous* spatial structure of the population influences the way its members interact, with interaction possible only within spatial neighbourhoods. The recurring theme of these studies is that when altruists cluster together they share the benefits of their behaviour mainly among themselves, and this can favour the spread of altruism. Our work is different in that we have a *discrete* spatial structure: individuals can migrate to different units, but the population itself within a unit is well-mixed; all its members interact in the same way with one another and everyone benefits from the mutants’ altruism in the same way. In this sense, our model is more similar to Refs. [30, 57, 75].

## 8 Parameter values for simulations in Figs. 1 and S1

*We detail the parameter values used to obtain the simulation results plotted in Figs. 1 and S1*

The parameter values for Figs. 1 and S1 are not derived from the scientific literature and are not meant to describe a realistic system. Rather, they are chosen to show the increase in mean mutant copy number and the increase in mean heteroplasmy with a limited computational effort in the simplest possible systems. Realistic parameter values for simulating skeletal muscle fibres models as a chain of units (Fig. 2) are reported in the next section.

### 8.1 Fig. 1

In Fig. 1C, All simulations are run for 7 · 10^2^ days. All systems have *μ* = 7 · 10^−2^/day, *c* = 10^−3^/day, with parameters that vary by panel specified below. All units are initialised (in steady state) with copy number (*w*_0_, *m*_0_) = 100, corresponding to an initial heteroplasmy *h*_0_ = 0.5. Averages have been taken over 2 · 10^3^ realisations. Error bars are SEM.

In the top-left subpanel, we simulate the two-unit deterministic system (Eq. (S36)), with the above parameters and *N_ss_* = 150, *δ* = 0.5, *γ* = 7 · 10^−4^/day. The black dots refer to the neutral system, the green dots refer to the system with preferential elimination of mutants (Eq. (S17)), with *ϵ* = 10^−3^*μ*. The top-right subpanel refers to the single-unit stochastic system (Eq. (S11)), with *N_ss_* = 150, *δ* = 0.5. The black dots refer to the neutral system, the green dots refer to the preferential elimination of mutants (Eq. (S17)), with *ϵ* = 10^−3^*μ*.

The bottom-left subpanel refers to the two-unit stochastic system (Eq. (S37)), with *N_ss_* = 200, *δ* = 1, *γ* = 3.5 · 10^−2^/day. The black dots refer to the neutral system, the green dots refer to the deterministic system with preferential elimination of mutants (Eq. (S17)), with *ϵ* = 10^−3^*μ*.

The bottom-right subpanel refers to the two-unit stochastic system (Eq. (S37)), with *N_ss_* = 150, *δ* = 0.5, *γ* = 3.5 · 10^−2^/day. The black dots refer to the neutral system, the green dots refer to the deterministic system with preferential elimination of mutants (Eq. (S17)), with *ϵ* = 10^−3^*μ*.

Fig. 1D refers to a chain of 340 units, with *μ* = 7·10^−2^/day, *c* = 0.4/day and diffusion parameterised by the hopping rate *γ* = 0.12/day. In the left subpanel, we simulate a chain of deterministic units with *δ* = 2*/*3. In the mid subpanel, we simulate a chain of neutral stochastic systems with *δ* = 1. In the right subpanel, we simulate a chain of neutral stochastic systems with *δ* = 2*/*3.

Fig. 1E refers to the same stochastic system as the right subpanel of Fig. 1D, except that preferential elimination of mutants (Eq. (S17)) has been added, with *ϵ* = 7.5 · 10^−2^*μ*.

### 8.2 Fig. S1

Fig. S1A depicts the steady state line (red) for a single-unit deterministic system (Eq. (S2)) with *N_ss_* = 500, *δ* = 0.5 (red line). In blue we depict a possible trajectory of the stochastic system (Eq. (S11)).

Fig. S1B refers to the single-unit system. Data is obtained simulating the deterministic ODE system Eq. (S2) (black lines) and the chemical reaction network Eq. (S11) via Gillespie algorithm (other lines) with parameters *μ* = 7 · 10^−2^/day, *δ* = 10^−1^, *c* = 2.5 · 10^−3^/day, *N_ss_* = 325. The system is initialised at steady state with *w*_0_ = 300, *m*_0_ = 250 corresponding to an initial heteroplasmy *h*_0_ ≈ 0.45. For the deterministic (black) and neutral stochastic (red) cases, mutants and wildtypes have the same degradation rate *μ*. For the cases of slow and fast preferential elimination, the degradation rate of mutants is increased by a constant *ϵ* (see Eq. (S17)). The increase in degradation rate is *ϵ* = 2.935 · 10^−4^/day for slow elimination (green) and *ϵ* = 1.1735 · 10^−3^/day for fast elimination (blue). For the deterministic system we plot mutant copy number *m*; for the stochastic systems we plot 〈*m*〉, averaging over an ensemble of 10 simulations. Error bars are standard error of the mean (SEM).

Fig. S1C is relative to the same system as panel B, but plotting (mean) heteroplasmy. For the deterministic system we plot heteroplasmy *h*; for the stochastic systems we plot 〈*h*〉, averaging over an ensemble of 10 simulations. Error bars are standard error of the mean (SEM).

Fig. S1D refers to the two-unit system. Data obtained simulating the deterministic ODE system Eq. (S36) (black line) and the chemical reaction network Eq. (S37) (other lines) with parameters *γ* = *μ* = 7 · 10^−2^/day, *δ* = 10^−1^, *c* = 10^−3^/day, *N_ss_* = 455. The chemical reaction network is simulated via the Gillespie algorithm. One unit is initialised with zero heteroplasmy *w*_10_ = 455, *m*_10_ = 0 and the other with *w*_20_ = 450, *m*_20_ = 50, corresponding to an initial heteroplasmy of 0.1. For the deterministic (black) and neutral stochastic (red) cases mutants and wildtypes have the same degradation rate *μ*. For the cases of slow and fast preferential elimination, the degradation rate of mutants is increased by a constant *ϵ* (see Eq. (S17)). The increase in degradation rate is *ϵ* = 1.5 · 10^−3^/day for slow elimination (green) and *ϵ* = 6.25 · 10^−3^/day for fast elimination (blue). For the deterministic system the quantity plotted is *h_mean_* = (*h*_1_ + *h*_2_)*/*2, the mean of the heteroplasmies of the two units. For the stochastic system, we plot (*h_mean_*) = ((*h*_1_ + *h*_2_))*/*2, averaging over an ensemble of 3.6 · 10^4^ simulations. Error bars are standard error of the mean (SEM).

## 9 Realistic estimates of parameter values and setup of numerical simulations

*We explain how realistic parameters have been estimated from independent experimental data, and report details on the large-scale stochastic simulations using these values*

In this section, we report and justify our best estimates of the parameter values used to simulate the spread of mtDNA deletions in the skeletal muscle fibres of rhesus monkeys. Specifically, we discuss the point estimates used to obtain Figs.2D, E, and the estimated ranges used to obtain Figs.2F. We derive these values from the scientific literature, in order to simulate a realistic travelling wave of mutants and compare the wave speed predicted by our simulations with that estimated from experiments (Fig. 2B).

Our model has a total of five parameters, all having an immediate and clear biological interpretation. We do not tune any of these parameters. Instead, we analyse experimental data to estimate parameter values that apply to skeletal muscle fibres of rhesus monkeys and use point estimates for our simulations (Figs. 2D, E). In Fig. 2D we show that stochastic simulations (Gillespie algorithm) model based on a net replicative advantage *k_RA_* for mutants predicts a travelling wave of heteroplasmy with a wave speed that is approximately 300 times faster than the observed speed. Crucially, in Fig. 2E we show that stochastic simulations of our model for the same system predict a wave of heteroplasmy with a speed of expansion comparable with observations. Setting up this simulation required some care, due to the slow dynamics caused by the large values of *N_ss_*. The wavefront of a travelling wave has the shape of a sigmoid whose steepness depends on its speed (see SI.14), therefore one needs to initialise the wavefront with the correct steepness such that the wavefront remains approximately stationary. For all the other simulations presented in the paper, this is not an issue since the heteroplasmy wavefront assumes the steepness corresponding to its constant speed in a short time transient (faster dynamics), that can then be neglected. Instead, for Fig. 2E we established an appropriate approximate initial steepness after a series of trials, by verifying that the shape did not change appreciably after running the simulation for a sufficient number of days (around 50). The chosen initial heteroplasmy profile is *h*_0_(*x*) = 1/(*e*^τ*x*−*b*^ + 1) for *x* > 0, with *τ* = 7.15 · 10^−4^/ μm and *b* = 3.915, over 500 units of length *L* = 30 μm. We also introduced approximations at the boundaries of the system. In order to simulate the effect of a macroscopic region of the muscle fibre taken over by mutants we fixed the value of the heteroplasmy of the leftmost unit of the chain at 1. At the right boundary, we truncated the values of heteroplasmy lower than 5 · 10^−3^ to zero, to rule out the possibility that edge effects alter the speed of expansion of the heteroplasmy wavefront. We verified that the rightmost units are mutant-free up to the longest simulated time.

In addition, in Fig. 2F we provide evidence that our conclusions are robust to the uncertainty in parameter values. We have drawn parameter values according to the distributions inferred from experimental data (see discussion below for details). We have then obtained distributions of the wave speeds predicted by a SSD and RA model (FK waves), inserting parameter values in the corresponding formulae for wave speed, namely Eq. (S43) with the appropriate interpretation of *k* for each case (setting *β* = 0 for FK waves). The approach used to obtain the distributions of the predicted wave speeds in is detailed in Ref. [103] and can be easily reproduced using the online tool at http://caladis.org/ and the parameter estimates given below. The distributions plotted in Fig. 2F confirm that SSD is strongly favoured to account for the expansion.

The parameters *μ, γ, c, N_ss_* are common to SSD and to the RA model, whereas the net mutant replicative advantage *k_RA_* is replaced by *δ* in our model. We estimate the degradation rate *μ* from the half-life of mitochondria. Reported values for the half-life in the literature range from a few to 100 days ^76,104–107^. We therefore assume that the half-life is uniformly distributed on the range (2,100) days. As a point estimate, we have chosen 10 days for the half-life, corresponding to a degradation rate of *μ* = 0.07/day.

*N_ss_* represents the average copy number per nucleus when only wildtypes are present. A study ^108^ reports *N_ss_* ~ 3500 for healthy human skeletal muscle fibres. We therefore suppose that for rhesus monkeys muscle fibres our uncertainty on *N_ss_* has a uniform distribution on the range (2000,5000). We have chosen *N_ss_* = 3000 since macaques are smaller animals. According to our empirical formula in Eq. (2), changing *N_ss_* from 3000 to 4000 would reduce the wave speed by about 10%, Therefore, we do not expect the uncertainty on *N_ss_* to affect our results significantly.

An important quantity in the study of muscle ageing, although not directly a parameter of our model, is the internuclear distance *L*. It has been reported that in skeletal muscle of healthy mice around 30 nuclei are present in 1*mm* of fibre length, i.e.~ 1 nucleus per 30 μm ^109^. We therefore take *L* = 30 μm as a point estimate, while assuming a uniform distribution on the range (25, 35) μm.

The hopping rate *γ* is linked to the diffusion coefficient of mtDNA molecules in the muscle fibre, through the relationship *D* = *γL*^2^ ^110^. Individual organelles in muscle fibres appear to be tightly packed. However, a study, Ref. [111], reported that the movement of nucleoids within organelles in these tissues can be effectively described as diffusion with a value of *D* corresponding to *γ* = 0.1/day. Therefore we choose this value as a point estimate of the individual mtDNA molecules. We quantify the uncertainty about the value of *γ* according to a uniform distribution on the range (0.05,0.15)/day The control strength *c* is the only parameter for which we do not have an optimal measurement. Our estimate is based on the fact that, in replicative cells, during a cell-cycle (≈ 1 day) the number of mtDNA roughly doubles. In this situation, in which a fast mitochondrial biogenesis is needed, we suppose that population is growing exponentially at a rate *λ*(*w* = 0, *m* = 0). For a doubling time of 1 day, this corresponds to

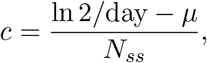

that for *N_ss_* = 3000 and *μ* = 0.07/day gives *c* = 2 · 10^−4^/day. Since *c* is not an independent parameter of our model, it is not present in the phenomenological wave speed formula Eq. (2).

The parameters *δ* and *k_RA_* correspond to the two ways of reproducing the increase in mutant carrying capacity (or density), respectively for our model and for a model based on a replicative advantage. From the experimental data ^21^ reported in Fig. 2C, in high-heteroplasmy regions density is approximately increased by a factor 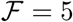 as reported also in Ref. [22]. We therefore choose 5 as the point estimate of 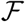 and we assume that 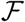 has a uniform distribution on the range (2,8). As for *δ*, we notice that in our model the carrying capacity *m*\* for mutants is obtained by setting in Eq. (S1) *λ* = *μ* and *w* = 0 gives, leading to *m*\* = *N_ss_/δ*. The carrying capacity for wildtypes is obtained by setting *λ* = *μ* and *m* = 0 in Eq. (S1), leading to *w*\* = *N_ss_*. Therefore, 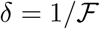 and we chose *δ* = 1/5 as a point estimate. In a replicative advantage model, the mutants have a replication rate given by Eq. (S1) (with *δ* = 1) and the addition of a net replicative advantage *k_RA_*, namely

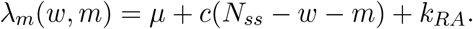

In order to reproduce an 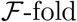 increase of mutant carrying capacity, it must be 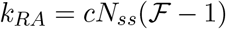. Indeed, setting *λ_m_* = *μ* and *w* = 0 yields *m*\* = *k_RA_/c + N_ss_*. Hence, we assume *k_RA_* to be uniformly distributed on the range (1, 7)*cN_ss_* and we choose *k* = 4*cN_ss_* as a point estimate, yielding

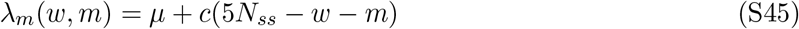

as the mutant replication rate in an RA model.

## 10 Revisited estimates of mutant loads in skeletal muscle fibres

*We provide new estimates and interpretation of mutant loads in skeletal muscle fibres, by analysing experimental data from muscle fibre biopsies*

Most existing modelling studies ^6,11,17^ for humans and rodents have tried to match predictions to the following end-of-life target mutant loads: 5 − 10% fibres must have reached a heteroplasmy level of ≥ 60%. We refer to the former quantity as *threshold cell-fraction* (TCF) and to the latter as *threshold heteroplasmy* (TH). The TH of 60% comes from a study on cybrid HeLa cells, stating that a 5196 base pair deletion needs to reach a cell-fraction of > 60% before the cell shows a reduction in COX activity ^112^. However, TH is cell-dependent^113^ and the 60% value for HeLa cells needs not be relevant for skeletal muscle fibres in humans and rodents. The TCF of 5% ^11,17^ is often justified on the basis of three studies ^114–116^. In the first study, 5% is the upper bound of the range (0.1-5)% ^114^. The second study found a maximum of 0.37% COX-negative fibres in human limb muscle over a large age range ^115^. The last study ^116^ did observe high percentages of COX-negative muscle fibres (~ 40%) but, it refers to a patient affected by a mitochondrial deletion disease, whereas we are interested in muscle ageing in healthy subjects.

We propose new estimates after examining data on skeletal muscle fibres of aged healthy mammals ^10,21,114,117^ and considering the spatial structure of muscle fibres. In these studies, hundreds of serial sections of length ~ 10*μ*m were cut along muscle fibres and stained for cytochrome *c* oxidase (COX) and succinate dehydrogenase (SDH) activities. Succinate dehydrogenase refers to complex II of the respiratory chain, the only complex fully encoded by the nucleus. Therefore, negative SDH activities indicate a nuclear defect, whereas normal or hyperactive activities *and* negative COX activities point towards mtDNA defects. An example of these data, from Ref. [21], is Fig. 2A, where additionally the heteroplasmy is measured for each section. Notice that, for simplicity, we have reported the spatial dimension as continuous, while in reality the measurements (the blue circles in the plot) refer to the section as a whole.

We suggest a TH of 90%, based on the fact that in Refs. [114, 117] regions of skeletal muscle fibres presenting mitochondrial dysfunctions were shown to contain > 90% of deletion mutations.

We now propose a new interpretation of the TCF. Previous mathematical studies modelled muscle fibres as a single units without spatial extension, with an unstructured and well-mixed mitochondrial population around a single nucleus. These models define TCF as the fraction of single-unit systems in which heteroplasmy exceeds TH. Having introduced a spatial model, we give an estimate of TCF by referring to experimental studies measuring heteroplasmy levels along the length of muscle fibres. In Ref. [21], of 10652 human vastus lateralis (VL) muscle fibres of aged (92 years old) subjects 98 fibres presented an abnormal region, approximately 1% of the total. In Ref. [10] 2115 muscle fibres of 34-year-old rhesus monkeys vastus lateralis (VL) were analysed, and 51 abnormal regions were found, corresponding to ≈ 2% of the total. These estimates for rhesus monkeys and humans refer to subjects at the end of their typical lifespans. We define TCF as the fraction of cells (i.e. fibres) that show a macroscopic abnormal region (where TH > 90%) and, on the basis or the above studies, we propose an end-of-life value of TCF≈ 1 − 2%.

## 11 Stochastic survival of the densest can account for mutational load in short-lived mammals

*We detail how we estimate the de novo mutation rate required by stochastic survival of the densest to reproduce the observed mutant loads in humans*

In the penultimate section of the main text we have claimed that, with respect to early neutral models of the clonal expansion of mtDNA deletions in skeletal muscle fibres, stochastic survival of the densest (SSD) dramatically lowers the value of the *de novo* mutation rate, *R_mut_*, required to yield a given mutant load. Here we explain this claim and provide a supporting mathematical argument.

Previous efforts to establish the ability of neutral genetic models to account for mammalian ageing have often focussed on understanding the compatibility between mutant loads and *R_mut_* through mathematical modelling. The approach is to develop a model predicting a certain amount of mutant molecules as a function of mutation rate, and then choose the *R_mut_* that matches experimentally observed (i.e. target) mutant loads. In Ref. [11] a neutral stochastic model of degradation and replication of mtDNA that used a mutation rate *R_mut_* = 5 × 10^−5^ per mtDNA replication, predicted that ≤ 4% of fibres will be COX-deficient due to high amount of mtDNA deletions by the 80th birthday of the subject, in agreement with experimental observations of healthy aged human muscle ^114,115^.

However, it was later argued that this mechanism cannot explain mutant loads for short-lived animals like rodents ^17^, which show similar accumulation patterns on much shorter time scales (≈ 3 years) ^4,54^. For the model to predict the observed mutant loads in short-lived animals such as mice and rats, *R_mut_* must be so high that it would produce an unrealistically high mutational diversity, whereas experimental studies report a single deletion that clonally expands ^20^. It has therefore often been assumed that neutral stochastic models cannot explain the mutational load in short-lived animals, and therefore models where mutants have a replicative advantage are needed.

In this work we have shown that SSD induces a wave-like clonal expansion of mutants without the need of a replicative advantage. In Fig. 4 we have plotted the reciprocal of the probability that a single founder mutation (arising in a muscle fibre modelled as a chain of units with *w* = *N_ss_* wildtype each), takes over the whole system (details in SI.12). The data are well described (*R*^2^ = 0.9995) by *a* + *bN_ss_*, with *b* = 0.301 ± 0.002, *a* = 1.5 ± 0.6 (linear fit). Based on this regression analysis, we estimate a fixation probability of *P_f_* = (9.47 0.06) 10^−4^ for *N_ss_* = 3500, a value plausible for humans. In the previous section SI.10 we referenced experimental studies^10,21,114,117^ reporting that in abnormal regions of muscle fibres, heteroplasmy is higher than 90%. In the following we will consider that abnormal regions have heteroplasmy 100%, i.e. that mutants have fixed. This approximation is not drastic and allows for a mathematically transparent argument.

The fraction of fibres that show an abnormal region – denoted by TCF, see previous section SI.10 – can be related to the number *M* of mutations that occur in a lifetime by the formula *MP_f_* = TCF. We assume that these *M* mutations arise early enough to have time to take over a macroscopic region of the system if not lost due to random degradation. In our model, a chain with *n* units each containing *N_ss_* molecules, if mtDNA mutates at a rate *R_mut_*, we expect to see *M* = *nN_ss_R_mut_*Δ*T* mutations arising in a time interval Δ*T* (measured in multiples of the characteristic time for a single mtDNA molecule to replicate). If we define that mutations occurring in the first Δ*T* of life have enough time to expand (if successful), we have

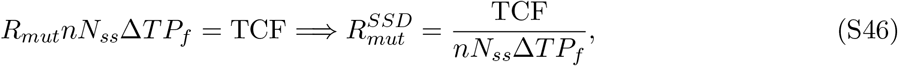

where by 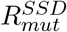 we denote the estimate of *R_mut_* based on SSD.

A conservative assumption is that mutations that spread to be observable are those arising in the first 10 years of life. Considering that, on average, a mtDNA molecule replicates every 10 days in a steady state situation (see estimate of *μ* in SI.9), 10 years correspond to 365 replications, i.e. Δ*T* = 365 replications. Typical skeletal muscle fibres are 3 − 12cm long, which means *n* ~ 1000 − 4000 (see estimate of *L* in SI.9). Inserting these estimates and *P_f_* = 9.47 10^−4^, *N_ss_* = 3500 in Eq. (S46), we obtain *R_mut_* = 4.1 10^−9^ 1.6 10^−7^ per replication, two-to-four orders of magnitude smaller than the estimate in the reference study ^11^.

The above argument is valid under the following assumptions:

1. A fibre can be divided into units, the building block of our model. A mutation can arise in a single unit and can expand into neighbouring ones.
2. An abnormal region is caused by a single mutation event. It might be that while a deletion is expanding, an *identical* one arises in the same region of the muscle fibre. We assume that this does not happen in our argument, as mutations are rare event and most of them are lost due to random degradation.
3. A mutation event cannot give rise to more than one abnormal region. Although it may be that distinct but nearby abnormal regions in a single fibre are the result of a single mutation event, this was not observed in the studies we investigated.

Neglecting spatial structure caused early models to miss the travelling wave of mutants driven by SSD and to require unrealistically high values of *R_mut_*. As SSD increases the probability that a single founder mutation takes over a long fibre, it also dramatically decreases the *de novo* mutation rate required (i.e. number of mutations) to yield observed mutant loads. More generally, what determines mutant load over time, and hence the possible progression of sarcopenia, is not only *R_mut_*, as assumed in early studies, but rather the speed of the wave of mutants. Different animals can have different mtDNA turnover rates, copy numbers, type of mutation and diffusion rate that, according to Eq. (2), affect the wave speed. Different species might thus be expected to experience sarcopenia on different time scales. In particular, according to our formula Eq. (2), short-lived animals are predicted to have higher wave speeds for a larger proportion of fibres, since they have more glycolytic, low-copy number fibres ^34^. For these two reasons – decreased *R_mut_* and possibly faster waves – SSD has the potential to account for sarcopenia also in short-lived animals.

## 12 Estimation of fixation probabilities in Fig. 4

*We explain the procedure used to estimate the fixation probabilities plotted in Fig. 4*

In Fig. 4 we have regressed the reciprocal of the fixation probabilities (of a founder mutation in a chain of units) as a function of the size of the steady-state wildtype population in which the founder mutation arises, which coincides with *N_ss_*. The analysis has shown a convincing linear relationship (*p* = 2 · 10^−19^, *R*^2^ = 0.9995).

We have estimated the fixation probabilities for 10 ≤ *N_ss_* ≤ 10^3^. For values 10 ≤ *N_ss_* ≤ 100, we have simulated a large ensemble of *N_sim_* ≈ 2.5 · 10^4^ muscle fibres, modelled as chains of hundreds of units, with a founder mutation in the middle of the chain. We have run the simulations until, in every fibre, the mutation had either died or taken over (*h* = 1) a macroscopic region of the fibre (~50-100 units). In the latter case, we have assumed that the mutants would have taken over the entire fibre with more time, excluding extremely unlikely fluctuations that could have led to the extinction of the entire mutant population (of size ~ 10^3^ here) occupying a large fraction of the fibre. The fixation probability *P_f_* has then been estimated as *N_mut_/N_sim_*, where *N_mut_* is the number of fibres mutants took over. In every case, we have ensured that at the boundaries of the chain were free or almost free of mutants, tolerating a heteroplasmy at the boundaries of at most 1/100.

For *N_ss_* ≥ 500, simulations with fibres long enough that mutants do not reach the boundaries are extremely slow. We have therefore estimated the fixation probability as shown in Fig. S3. We have run a large number *N_sim_* of simulations (*N_sim_* ≈ 25000) with a founder mutation at time 0, and recorded the number of simulations where mutants are present over time. We noticed that this number decreases exponentially after an initial transient (not shown in the plots) of ≈ 4000 days in which it decreases drastically (from ≈ 25000 to ≈ 150). Fitting an exponential decay to the data allows us to estimate the height of the horizontal asymptote of the curve, namely the stationary value of the number of simulations with surviving mutants. Since this asymptotic value coincides with *N_mut_*, we use it to estimate *P_f_*, again as *N_mut_/N_sim_*. As above, we are ignoring extremely rare events, where a large number of mutants (~ 10^4^ here), that have taken over a macroscopic fraction of the fibre (hundreds of units here), would go extinct. We stopped the simulations when running them for longer, and adding data for longer times to the exponential fits in Fig. S3, changed the estimate of *N_mut_* – i.e. the horizontal asymptotes – by less than one unit. More detail on the regression analysis are in SI.16.

We estimated the standard deviation *σ* of *P_f_* modelling the phenomenon as a binomial experiment, namely as *σ* = [*P_f_* (1−*P_f_*)*/N_sim_*]^1/2^. In Fig. 4 we plotted the reciprocal of the fixation probabilities, i.e. 1*/P_f_* = *N_sim_/N_mut_*. For the uncertainties on the values of 1*/P_f_*, which are the error bars plotted in Fig. 4, we used for each value of *N_ss_* the standard deviation of the random variables *Y = Nss/X*, where *X* ~ Binomial(*N_ss_, P_f_*). We estimated this standard deviation simulating 10^7^ samples of *Y* for each value of *N_ss_*.

In Table S1, we list the values of *N_mut_, N_sim_* and the corresponding estimates for each value of *N_ss_* used.

## 13 Quadratic and reciprocal feedback produce a wave of heteroplasmy

*We present variants of the replication rate in Eq. (1), that also produce stochastic survival of the densest, a mechanism that is therefore independent of the mathematical details of our model*

In Fig. S4C, D we show that other types of feedback controls produce an analogous travelling wave of heteroplasmy without the need of a replicative advantage for mutants. The two additional types of controls are the *quadratic* control, namely

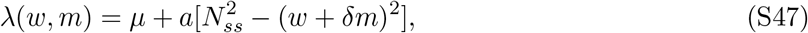

and the *reciprocal* control, given by

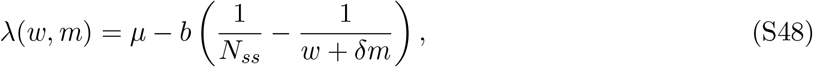

where *a* and *b* are control strength parameters. Importantly, these control mechanisms all produce wave speeds that decrease when *N_ss_* increases, a fundamental difference between our model and models based on a net replicative advantage for mutants.

The data reported in Fig. S4C, D are obtained averaging over 400 realizations with the following parameters. For quadratic feedback, Fig. S4C and Eq. (S47): *μ* = 7 · 10^−2^/day, *γ* = 0.12/day, *δ* = 2/3, *a* = 0.4/day, *N_ss_* = 4. For reciprocal feedback, Fig. S4D and Eq. (S48): *μ* = 7·10^−2^/day, *γ* = 10^−2^/day, *δ* = 0.1, *b* = 1.5/day, *N_ss_* = 3. The small values of *N_ss_* are chosen for numerical convenience, since a smaller copy number produces a faster dynamics and faster waves. In the light of these results, we argue that the mechanism of stochastic survival of the densest is independent of the mathematical details of our model, provided that the three key factors of density difference, stochasticity and spatial structure (with diffusion) are present.

## 14 Relationship between steepness and speed of a travelling wave

*We provide computational evidence that the relationship between steepness and speed of a travelling wave driven by stochastic survival of the densest is the same as Fisher-Kolmogorov waves*

This section is based on Secs. 13.2 and 13.3 of Ref. [53]. Let us consider the classical Fisher-Kolmogorov reaction-diffusion equation

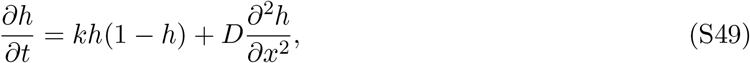

where *h* is heteroplasmy, *k* is the reaction rate and *D* is the diffusion coefficient. This equation admits travelling wave solutions moving in the positive *x*-direction with speed *c* given by

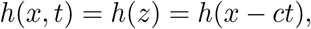

having defined *z* = *x* − *ct*. If the mutants of our system had a replicative advantage *k*, Eq. (S49) would describe the wave-like expansion of the high-heteroplasmy front in the continuous-space approximation. If the system is initialised such that

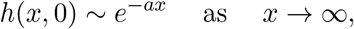

for arbitrary *a* > 0, the heteroplasmy wavefront will evolve into an approximately sigmoidal shape with a steepness *τ* linked to the wave speed through

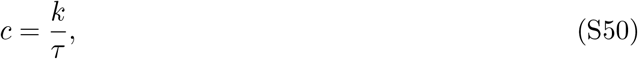

meaning that the slower the wave, the steeper the wavefront (larger *τ*).

We speculate that waves driven by stochastic survival of the densest can be described by an equation analogous to Eq. (S49), with an effective reaction rate induced by stochastic survival of the densest, that we hypothesise is given by Eq. (S44). Therefore, also for our model we expect that steeper waves will be slower. In absence of a formal proof, we have verified this intuition computationally. In Fig. S5 we plot the travelling waves of heteroplasmy for two systems that differ only in copy number *N_ss_*. The blue curve corresponds to a system with a smaller *N_ss_* and hence, according to Eq. (2) presenting a faster wave; the orange curve corresponds to a larger *N_ss_* and hence a slower wave. As expected, the slower wave has a steeper wavefront.

In Fig. 3 we have reported experimental data on human (panels A and B) and rat (panels C and D) muscle fibres, showing that the steeper wavefront corresponds to the fibre with a larger *N_ss_*. We can conclude that the steeper wavefront corresponds to a slower wave. Hence, the data reported in Fig. 3 support the prediction of our model that slower waves travel in fibres with a larger *N_ss_*.

The data plotted in Fig. S5 are obtained simulating a chain of 550 units evolving under linear feedback control (Eq. (1)), with parameters *μ* = 7 · 10^−2^/day, *c* = 0.4/day, *γ* = 0.12/day, *δ* = 2/3. The faster (flatter) blue curve corresponds to *N_ss_* = 6, while the slower (steeper) orange one to *N_ss_* = 2.

## 15 Rhesus monkeys data collection

*We explain how we obtained the data points plotted in Fig. 2B from experimental data*

The data plotted in Fig. 2B on the length of the abnormal regions in muscle fibres of rhesus monkeys was originally published in Ref. [10]. The measurements were performed on skeletal muscle tissues of 11 different animals and the data set consists of the lengths of several abnormal regions for each subject. For each animal, we only use the length of the longest abnormal region, that we assume has originated from a mutation at birth. We consider shorter regions to be the results of mutations that have arisen later in life. Although these assumption are unlikely to be precisely true, our analysis is enough to give an indicative order-of-magnitude estimate, that can be considered as a moderately tight lower bound on the speed of expansion.

## 16 Statistical methods

*We provide information on all the statistical analyses peformed*

Error bars in Figs. 1C and S1B, C, D are standard error of the mean (SEM). Error bars in Fig. 4 are standard deviations, obtained as explained in SI.12.

In Fig. 2A, the functional form fitted is a logistic function (sigmoid) 1/(1 + *e^τ^ ^x^*). The variable *x* is the position along the muscle fibre, centred at the point where *h* = 1. The parameter is *τ* = (0.25 ± 0.01)/μm. The values are obtained using non-linear least squares, and the uncertainty is the standard error. In Fig. 2B there are 11 points. With ordinary least squares we obtain *R*^2^ = 0.758, *p* = 5 · 10^−4^, *t* = 5.3 for a line of equation *y* = *a* + *bx*, with *a* = (−482 ± 229) μm and *b* = 0.131 ± 0.025 μm/day (uncertainty is standard error). The wave speeds relative to Figs. 2D, E and those corresponding to the points in Fig. S2A have been calculated as explained in SI.5. In Fig. 2F 9 · 10^6^ values have been drawn for each of the three histograms.

For the fits in Figs. 3A, C non-linear least squares is used. The values obtained are reported in the caption of Fig. 3, and the uncertainty is standard error. The statistical test performed on the data in Figs. 3B, D is a one-tailed Welch’s test (unequal-variance *t*-test). For data in Fig. 3B, the two samples are of size 15 and 7, with measurements taken from different samples, meaning distinct slices of the same muscle fibres; means for copy numbers are 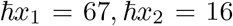 standard deviations *σ*_1_ = 39, *σ*_2_ = 6.5. Welch’s test gives *p* = ·10^−4^, *t* = 4.69, *d* = 1.48, where the effect size *d* is the unequal-variance Cohen’s *d*. For data in Fig. 3B, the two samples are of size 6 and 5, with measurements taken from different samples, meaning distinct slices of the same muscle fibres; means for copy numbers are 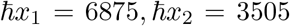 standard deviations *σ*_1_ = 3840, *σ*_2_ = 1700. Welch’s test gives *p* = ·0.6, *t* = 1.76, *d* = 1.00 (Cohen’s *d*). The elements of the box plots are minimum value (lower whisker), first quartile, median, third quartile and maximum value (upper whisker).

In Fig. 4, we have 13 points, with error bars being standard deviations, obtained as explained in SI.12. By linear least squares, we fit a line of equation *a* + *bN_ss_*, with *b* = 0.301 ± 0.002, *a* = 1.5 ± 0.6, with *p* = 10^−20^ (for *b*) and effect size *R*^2^ = 0.9995 (square of the Pearson’s correlation coefficient *r*). The uncertainty reported is standard error.

In Fig. S1, data from numerical simulations are plotted. Error bars are SEM and averages are taken over a sample of 10^4^ simulations. Details in SI.8.

In Fig. S2A, there are 110 points, and maximum likelihood estimation is used to fit the expression 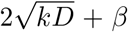 obtaining *β* = 0.207 ± 0.007, with effect size *R*^2^ = 0.99. Uncertainty reported is the standard error. Notice that the variable *k* is defined in the main text and in SI.5 and *D* is the diffusion coefficient, therefore *β* is the only parameter estimated.

In Fig. S3, non-linear least squares was used, fitting the function *y = ae*^−*bt*^ + *c*. For *N_ss_* = 500, the estimated values are *a* = 21.8 ± 0.1, *b* = (4.7 ± 0.1) · 10^−4^/day, *c* = 150.1 ± 0.1. For *N_ss_* = 1000, the estimated values are *a* = 55.8 ± 0.3, *b* = (3.37 ± 0.06) · 10^−4^/day, *c* = 85.6 ± 0.4. Uncertainty reported is standard deviation. See SI.12 for more details on how the synthetic data was obtained.

In Fig. S4, synthetic data is plotted, averaged over an ensemble of 400 simulations; more details in SI.13. In Fig. S5, the average is over 4000 simulations; more details in SI.14.

## 17 Author contributions

FI observed and characterised the noise-induced waves, performed analytical calculations and statistical analysis, coded the model and performed the simulations, interpreted the results, obtained new estimates of the *de novo* mutation rate, drafted the paper and supplement. HH first investigated spatially structured systems, observed and interpreted early evidence of stochastic survival of the densest and revisited the estimates of mutant loads in muscle fibres. JA supported FI with mathematical techniques, critically revised the manuscript and significantly contributed to the presentation of the results. NJ formulated and designed the research, interpreted the results, supervised, and contributed to, the analytics, statistics and computation and edited the drafts.

## 18 Data Availability Statement

No further experiment was performed for this work, previously published data was analysed and the relevant publications were referenced in the paper. For simplicity, we report the references here. Data in Fig. 2A, C was published in Ref. [21]. Data in Fig. 2B was published in Ref. [10]. Fig. 2A, B is from Ref. [3], Fig. 2C, D is from Ref. [4]. Data from simulations can be reproduced by the provided code (SI.19. The authors welcome questions about simulations and requests of simulated data.

## 19 Code availability

The C code for the linear feedback control model is available at https://github.com/ferdinando17/Survival-of-the-Densest-Code. The authors welcome questions. The parameter values currently specified in the code allow the reproduction of Fig. S1.D, the simplest system in which survival of the densest is observed. Other plots can be reproduced with the parameter and instructions specified in this work (SI.8, SI.9). Notice that in order to obtains plots such as Fig. 4 and Fig. 2.E we employed hundreds processors running in parallel for several months.

---

**Figure S1:**
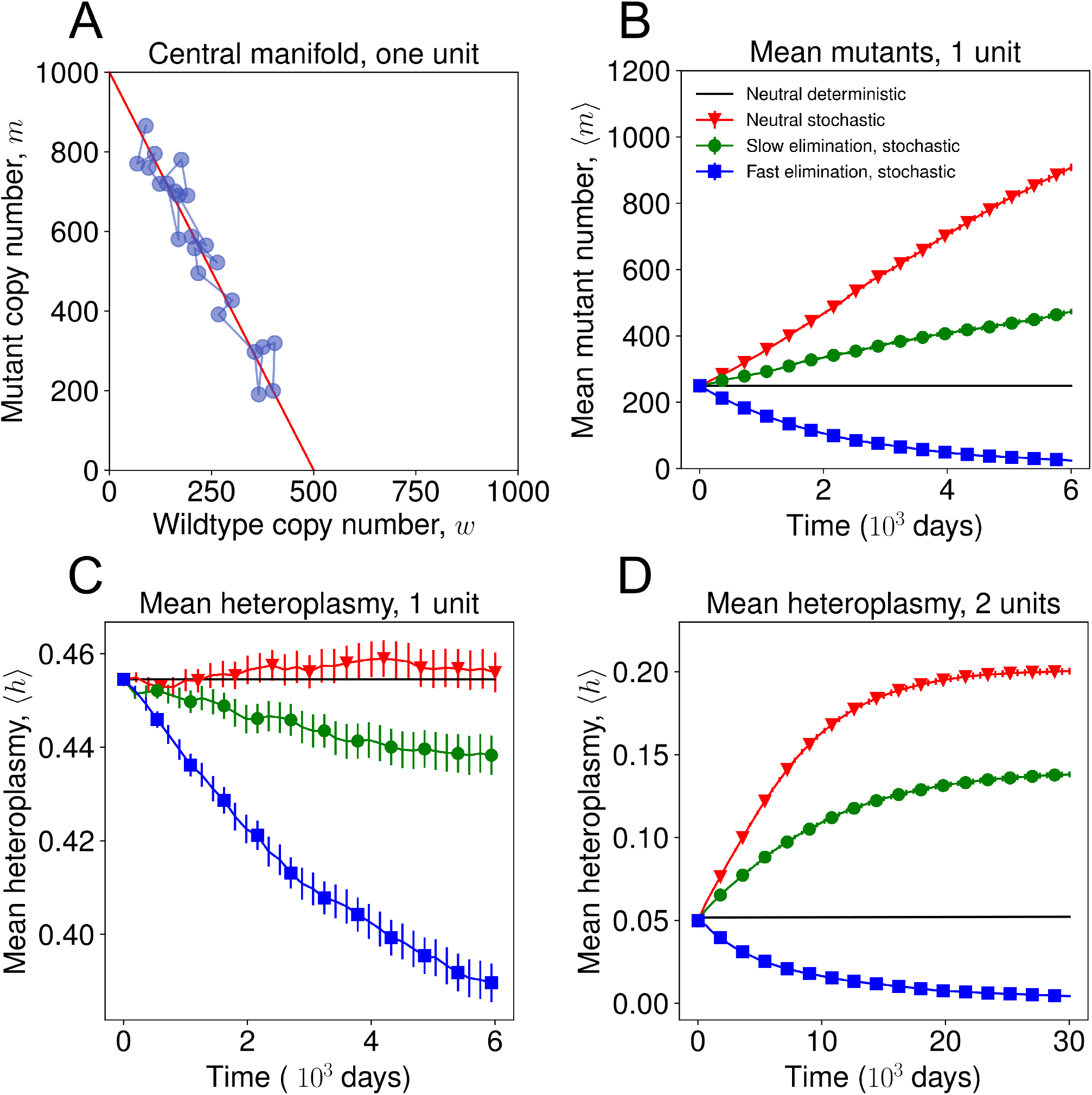
(A) The single-unit system fluctuates around the steady state line (red line), on average moving toward regions of higher *m* (average proportion of mutant (*h*) stays constant). In the plot, *N_ss_* = 500 and *δ* = 0.5. (B) In the stochastic single-unit system, mean mutant copy number increases in the neutral model (red) and in presence of weak preferential elimination of mutants (green). This is a consequence of higher carrying capacity for mutants (*δ* < 1) and noise. In the corresponding deterministic neutral model, mutant copy number remains constant (black). Parameter values in SI.8. Error bars are standard error of the mean (SEM). (C) In the single-unit system, heteroplasmy is constant in the neutral deterministic system, as well as mean heteroplasmy in the neutral stochastic system. Any selective elimination of mutants in the stochastic system causes a decrease in mean heteroplasmy, while mean number of mutants can still increase (green line in panel B). Parameter values in SI.8. Error bars are SEM. (D) In the stochastic two-unit system, mean mutant fraction (heteroplasmy) increases in the neutral model (red) and even if mutants are preferentially eliminated (green). Parameter values in SI.8. Error bars are SEM.

**Figure S2:**
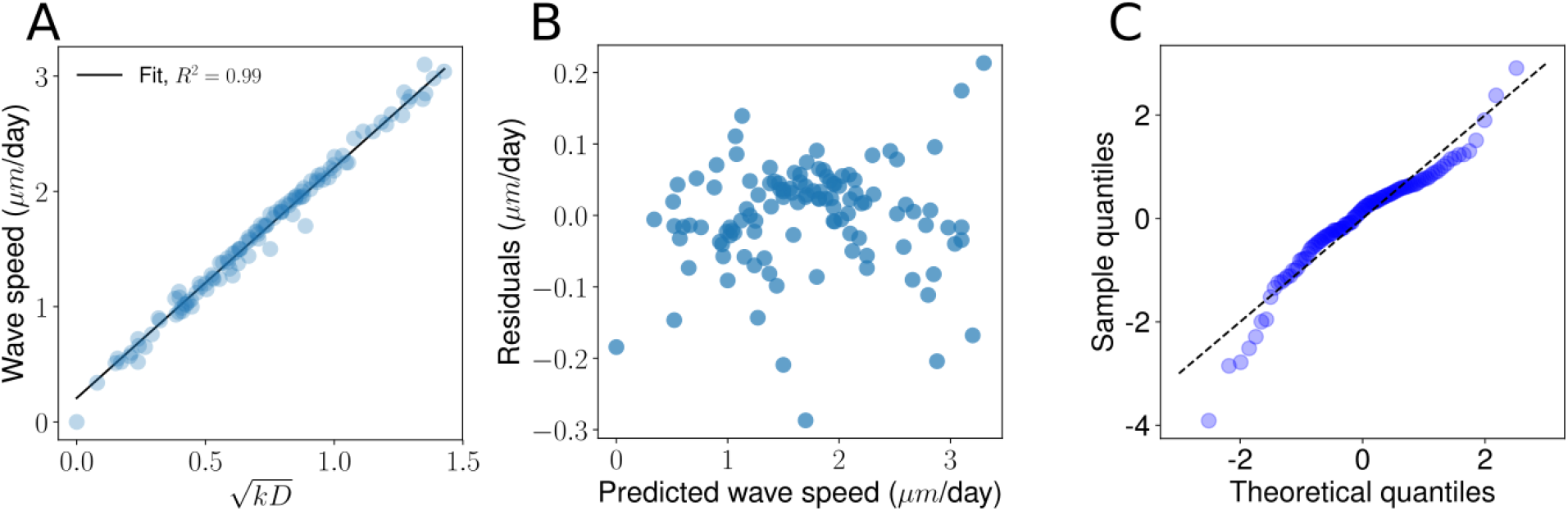
An empirical formula accurately predicts the survival of the densest wave speed obtained through stochastic simulations for a wide range of parameter values. A) The blue dots are 110 values of wave speed measured through stochastic simulations of the linear feedback control for varying values of the parameters *N_ss_, μ, γ* and 0 < *δ* < 1. The black line is the linear fit against the variable 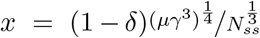 The excellent fit (*R*^2^ = 0.99) shows that the wave speed *v* is well predicted by Eq. (S43), except for *x* → 0. B) Although the residuals do not follow a Gaussian distribution (Shapiro-Wilk, *p* < 10^−4^), the absence of a clear pattern in the plot suggests that Eq. (S43) is a satisfactory approximation in the parameter range considered. C) The Q-Q plot of the residuals against a normal distribution confirms the absence of a clear pattern.

**Figure S3:**
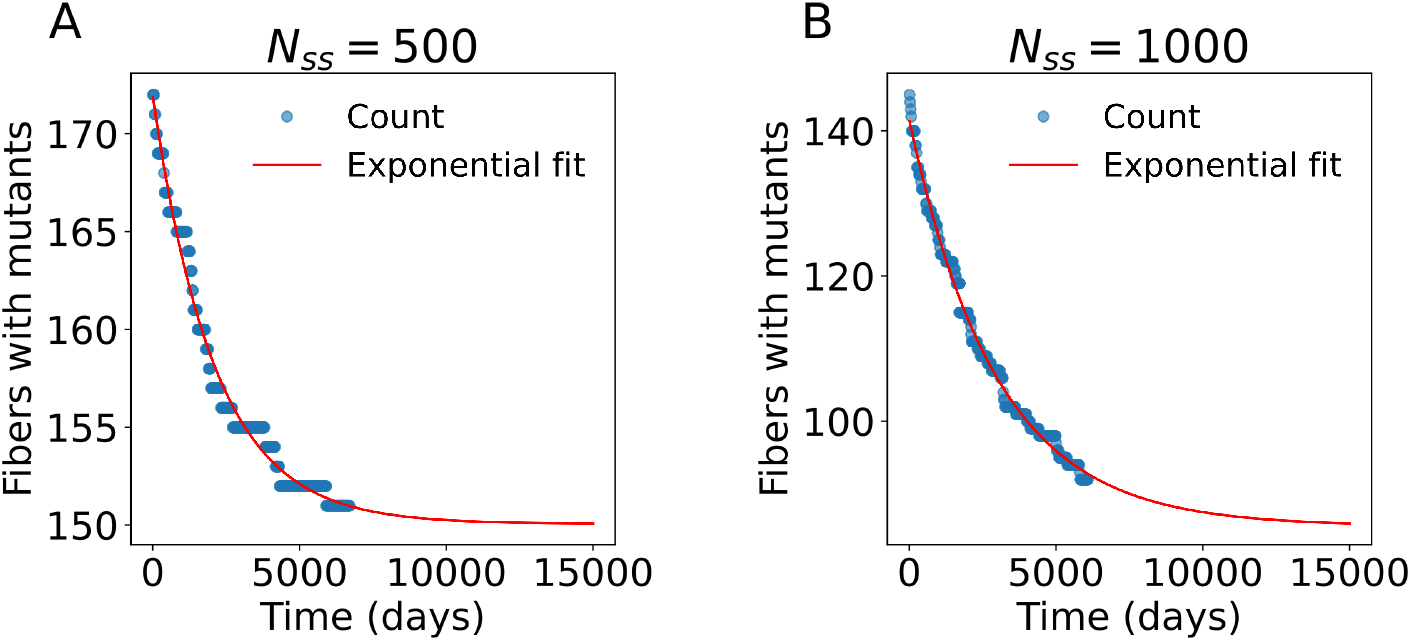
The above exponential fits allow us to estimate the fixation probability *P_f_* of a founder mutation in a chain of 500 (A) and 1000 (B) units. We have initialised an ensemble of *N_sim_* ≈ 25000 fibres with a founder mutation. Fitting an exponential to the points, we have estimated the stationary value *N_mut_* of the number of simulations with surviving mutants, that coincides with the number of simulations in which mutants take over the fibre (neglecting extremely unlikely extinction events). We then estimate *P_f_ = N_mut_/N_sim_*.

**Figure S4:**
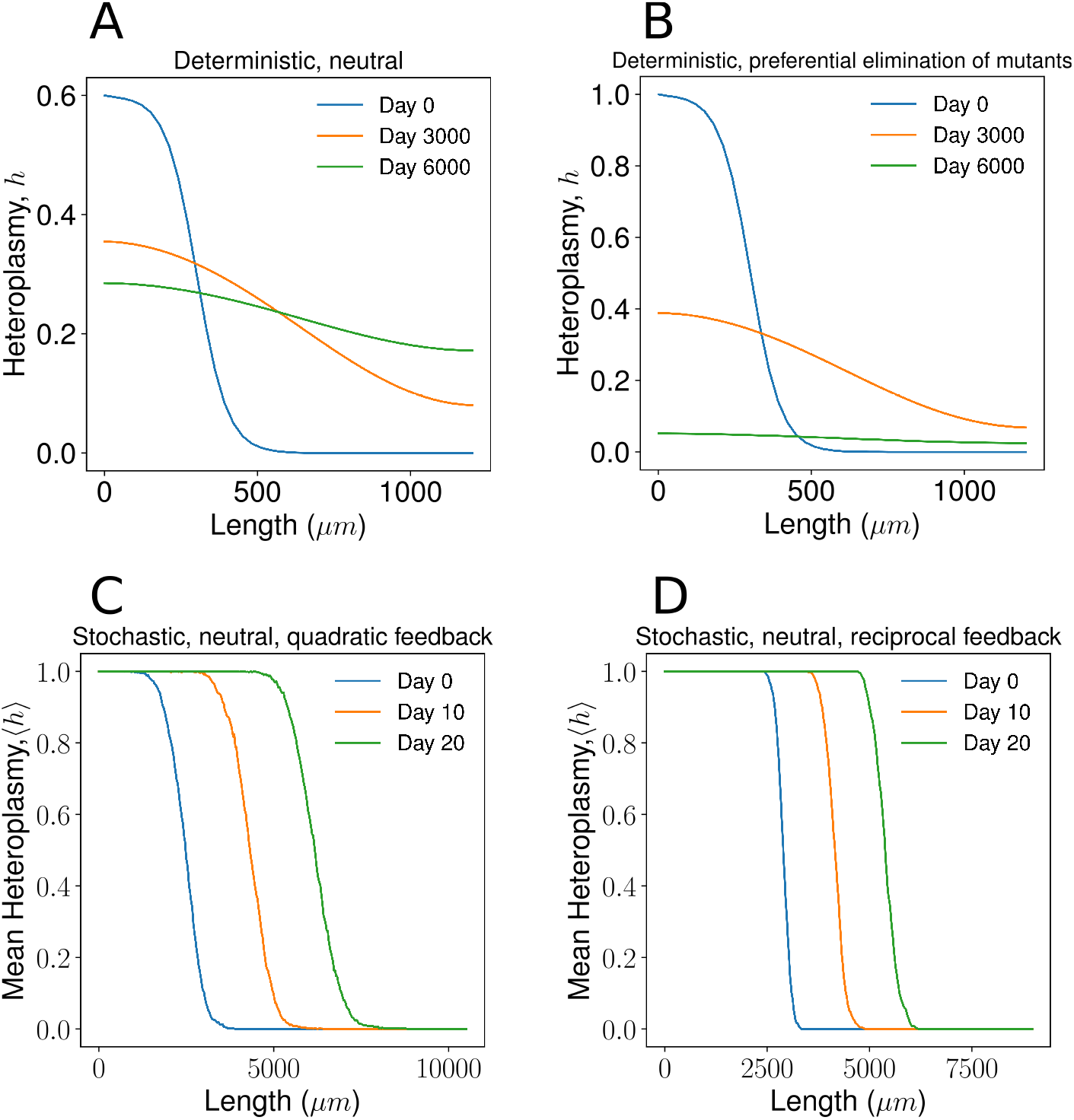
A deterministic model deployed on a chain does not produce mutant expansion despite differences in carrying capacity, while different choices of feedback control still produce a wave of mutants when stochasticity is present. (A) The deterministic ODE model of Eq. (S36) applied to a chain of units does not produce a travelling wave of heteroplasmy, even with larger carrying capacity for mutants (*δ* = 0.2). Rather, we observe diffusion of the heteroplasmy front. The curves are the heteroplasmy values obtained by ODE integration of a system analogous to Eq. (S36) for 41 units. (B) In the deterministic chain, when the degradation rate of mutants is higher than that of wildtype (1% in the plot), mutants go extinct despite a higher carrying capacity (*δ* = 0.2). (C) Quadratic and (D) reciprocal feedback control (see SI.13) produce a qualitatively similar wave-like expansion of mutants in a stochastic model.

**Figure S5:**
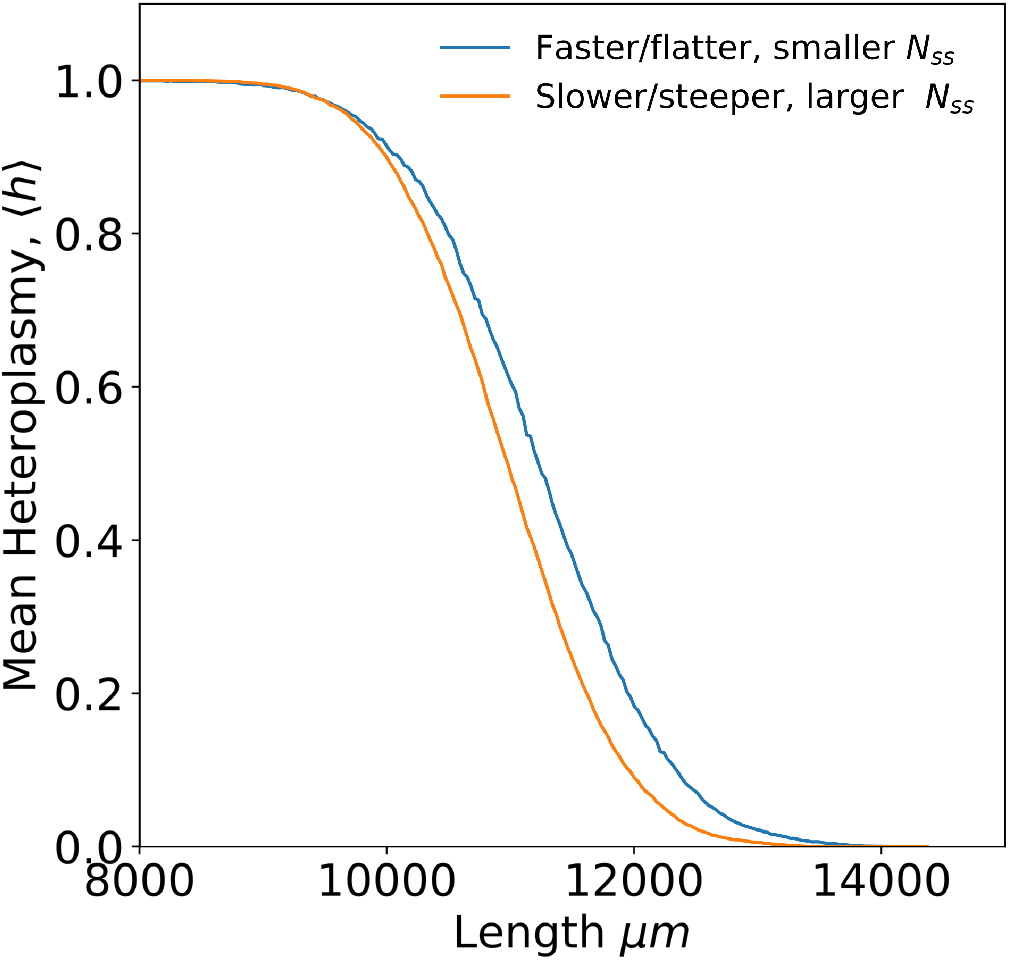
The two curves are the travelling wavefronts of two systems that differ only in the average wildtype copy number *N_ss_*. The smaller the *N_ss_*, the faster the wave, according to Eqs. (S43) and (S44). The blue curve is the wavefront propagating in the system with the smaller 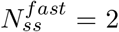, while the blue curve is relative to 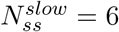. The plot shows that faster waves have a flatter wavefront, as it is the case for FK waves. Average over an ensemble of 4000 simulations.

**Table S1:**
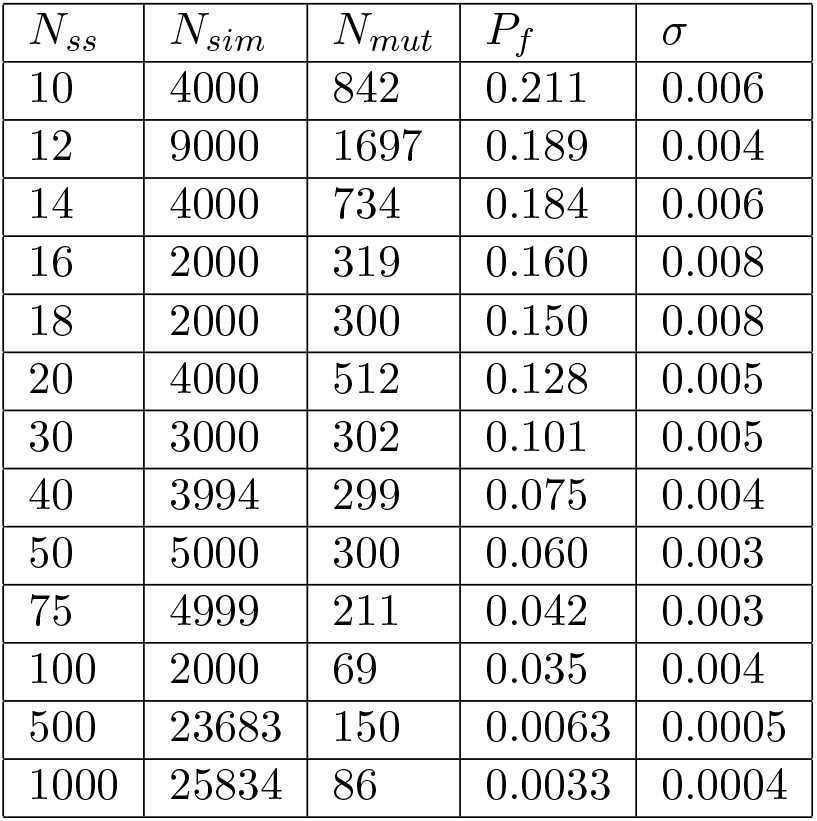
Estimates of *P_f_* and standard deviation *σ*, with corresponding values of *N_mut_, N_sim_*, for different values of *N_ss_*

## Notes

### Competing Interest Statement

The authors have declared no competing interest.

### Summary of Updates

The main text has been shortened and rebalanced. New results have been added, namely the fixation probability of a founder deletion mutation in muscle fibres and an estimate of the mutation rate required to reproduce experimentally observed mutant loads. Figures 1,2 and 4 have been updated. Figure 1 now clarifies that survival of the densest is an intrinsically stochastic effect. Supplement has been simplified and re-organised. Code has been provided (GitHub repository).

