## Supplementary material for "Stochastic survival of the densest and mitochondrial DNA clonal expansion in ageing": Mathematica notebook for stochastic dimensionality reduction.

### Derivation of effective SDE for mutant dynamics

```
ClearAll["Global`*"]
$Assumptions = z ∈ Reals && Nss ∈ Reals && c ∈ Reals && δ ∈ Reals && c > 0 && δ > 0 && Nss > 0;
```

### Definition of system of coupled SDEs

```
f = {c * m * (Nss - w - δ * m), c * w * (Nss - w - δ * m)};

G = {{ Sqrt[m * (c * (Nss - w - δ * m) + 2 * μ)], 0}, {0, Sqrt[w * (c * (Nss - w - δ * m) + 2 * μ)]}};
x = {m, w};
```

- Computing Hessians

```
H1 = D[f[[1]], {{m, w}, 2}];
H2 = D[f[[2]], {{m, w}, 2}];
```

- Defining variable constrained to central manifold (CM), with z=m (mutant copy number).

```
γ = {z, Nss - δ * z};

γ' = Simplify[D[γ, z]];
```

#### Definition of Jacobian. Evaluation on central manifold  (CM) and eigendecomposition.

```
J = D[f, {x}];

w = Nss - δ * m;

v = Eigenvectors[Transpose[J]][[1]];

NormalizationFactor = Simplify[Dot[v, γ']];

P = v / NormalizationFactor;
```

Printed by Wolfram Mathematica Student Edition

```
W = Transpose[Eigenvectors[J]];
```

### Computation of pseudo-inverse of Jacobian, of projection matrix P and of matrix Q_1

```
Jpi = FullSimplify[W.DiagonalMatrix[{0, 1/Eigenvalues[J][[2]]}].Inverse[W]];

Pf = FullSimplify[IdentityMatrix[2] - Jpi.J];

Q1 = -FullSimplify[Jpi[[1, 1]] * Transpose[Pf].H1.Pf +
     Pf[[1, 1]] * (Transpose[Jpi].H1.Pf + Transpose[Pf].H1.Jpi) +
     Jpi[[1, 2]] * Transpose[Pf].H2.Pf +
     Pf[[1, 2]] * (Transpose[Jpi].H2.Pf + Transpose[Pf].H2.Jpi)];
```

### Computation of drift term

```
Drift = FullSimplify[Tr[G.Transpose[G].Q1]/2]
```

$$\frac{2\,m\,(-1+\delta)\,(-Nss+m\,\delta)\,\mu}{Nss^2}$$

### Computation of diffusion term (the term that multiplies the Wiener process)

```
Eta = {{η1, η2}} ; (*Vector of Wiener noises*)
NoiseTerm = P.G.Transpose[Eta]
```

$$\left\{\frac{\sqrt{2}\,(Nss-m\,\delta)\,\eta 1\,\sqrt{m\,\mu}}{Nss}-\frac{\sqrt{2}\,m\,\eta 2\,\sqrt{(Nss-m\,\delta)\,\mu}}{Nss}\right\}$$

The above diffusion terms can be combined into a single one:

```
Var = FullSimplify[(P.G).(P.G)]
```

$$\frac{2\,m\,(-Nss+m\,(-1+\delta))\,(-Nss+m\,\delta)\,\mu}{Nss^2}$$

This is the variance of the Wiener process that drives the SDE.
